# Towards universal synthetic heterotrophy using a metabolic coordinator

**DOI:** 10.1101/2022.11.02.514957

**Authors:** Sean F. Sullivan, Anuj Shetty, Tharun Bharadwaj, Naveen Krishna, Vikas D. Trivedi, Venkatesh Endalur Gopinarayanan, Todd C. Chappell, Daniel M. Sellers, Pravin Kumar R., Nikhil U. Nair

**Affiliations:** Department of Chemical & Biological Engineering, Tufts University, Medford, MA 02155; Kcat Enzymatic Private Limited, Bengaluru, Karnataka, India 560005; Department of Structural Biology and Center for Data Driven Discovery, St. Jude Children’s Research Hospital, Memphis, TN

**Keywords:** Gene regulation, sustainability, synthetic biology, metabolic engineering, substrate utilization, molecular dynamics

## Abstract

Engineering the utilization of non-native substrates, or synthetic heterotrophy, in proven industrial microbes such as *Saccharomyces cerevisiae* represents an opportunity to valorize plentiful and renewable sources of carbon and energy as potential inputs to biotechnological processes. We previously demonstrated that activation of the galactose (GAL) regulon, a regulatory structure used by this yeast to coordinate substrate utilization with biomass formation during growth on galactose, during growth on the non-native substrate xylose results in a vastly altered gene expression profile and faster growth compared with constitutive overexpression of the same heterologous catabolic pathway. However, this effort involved the creation of a xylose-inducible variant of Gal3p (Gal3p^S2514^^4^.^1^), the sensor protein of the GAL regulon, preventing this semi-synthetic regulon approach from being easily adapted to additional non-native substrates. Here, we report the construction of a variant Gal3pMC (metabolic coordinator) that exhibits robust GAL regulon activation in the presence of structurally diverse substrates and recapitulates the dynamics of the native system. Multiple molecular modeling studies confirm that Gal3p^MC^ occupies conformational states corresponding to galactose-bound Gal3p in an inducer-independent manner. Using Gal3p^MC^ to test a regulon approach to the assimilation of the non-native lignocellulosic sugars xylose, arabinose, and cellobiose yields higher growth rates and final cell densities when compared with a constitutive overexpression of the same set of catabolic genes. The subsequent demonstration of rapid and complete co-utilization of all three non-native substrates suggests that Gal3p^MC^-mediated dynamic global gene expression changes by GAL regulon activation may be universally beneficial for engineering synthetic heterotrophy.

## INTRODUCTION

The adoption of abundant and renewable substrates as inputs for biotechnology will be essential for creating a sustainable, circular bioeconomy ^1, 2^. But the inability of industrially important microbes, like *Saccharomyces cerevisiae* (henceforth, yeast), to utilize many potential substrates poses a major hurdle to realizing this goal. The existing paradigm for engineering the assimilation of non-natives substrates (i.e., synthetic heterotrophy) in yeast begins with the identification and constitutive overexpression of catabolic genes that enable the substrate to enter central carbon metabolism (CCM) where it is expected to be transformed into the key primary metabolites used for growth and biosynthesis. However, this method ignores the tight regulation of CCM and how these resources are distributed to accomplish cellular objectives ^3^. The consequences of this oversight are reflected in the need for subsequent interventions, including flux balancing ^4–6^, functional genomics ^7–11^, and adaptive lab evolution ^12–14^, undertaken to improve growth and/or substrate utilization rate by resolving conflicts with cellular processes. Moreover, existing efforts are often focused on a specific substrate or small set of substrates – a siloed approach that makes it hard to translate findings to other substrates and even disincentivizes holistic thinking about the limits of metabolic plasticity/adaptability in this yeast. This has, at least in part, motivated domestication of other yeasts with broader substrate ranges (e.g., *Scheffersomyces stipitis, Kluyveromyces marxianus*) for biomanufacturing applications.

An alternate approach that we advocate here, is that instead of combating natural regulation we should leverage existing regulatory structures that have evolved to coordinate complex phenotypes like substrate utilization with biomass formation and metabolite synthesis. One such system is the galactose (GAL) regulon that yeast uses to coordinate substrate catabolism with global metabolism during growth on the native substrate galactose. In this system, the interaction of galactose with sensor protein Gal3p enables it to relieve the repression of Gal80p on the transcription factor Gal4p via protein-protein interactions ^15^. Once freed from repression, Gal4p – the master activator of the GAL regulon – binds to its cognate Upstream Activating Sequences (UAS) to directly induce transcription of galactose catabolic genes (Leloir pathway) and indirectly modulates the transcript and/or protein abundance of numerous growth-associated genes (**Figure 1A**) ^16, 17^. We previously reported the construction of a variant of sensor protein Gal3p (Gal3p^Syn4.1^) that strongly activates the GAL regulon by the pentose sugars xylose and arabinose ^18, 19^. By placing heterologous genes for either xylose or arabinose catabolism under the control of GAL-responsive (Leloir pathway) promoters, the GAL regulon could be adapted for growth on these non-native substrates. Upon comparing this regulon-coordinated (REG) approach to growth with simple constitutive overexpression of the same catabolic genes, which we term a constitutive (CONS) approach, we found that our semi-synthetic regulon yielded superior growth rates on both xylose (µ = 0.24 h^−1^ vs. 0.11 h^−1^) and arabinose (µ = 0.27 h^−1^ vs. 0.06 h^−1^), respectively with minimal metabolic engineering ^19^. Significantly, a transcriptomic comparison showed relative upregulation of genes associated with cell division and mitochondrial biogenesis in the REG strain while the CONS strain showed upregulation of stress response and starvation-associated genes ^18^. This suggests that the GAL regulon dynamically reshapes the cellular response for growth through direct and/or indirect action of Gal3p-Gal80p-Gal4p to potentiate the cells for rapid growth ^17, 20^, irrespective of the identity of the available substrate.

**Figure 1:**
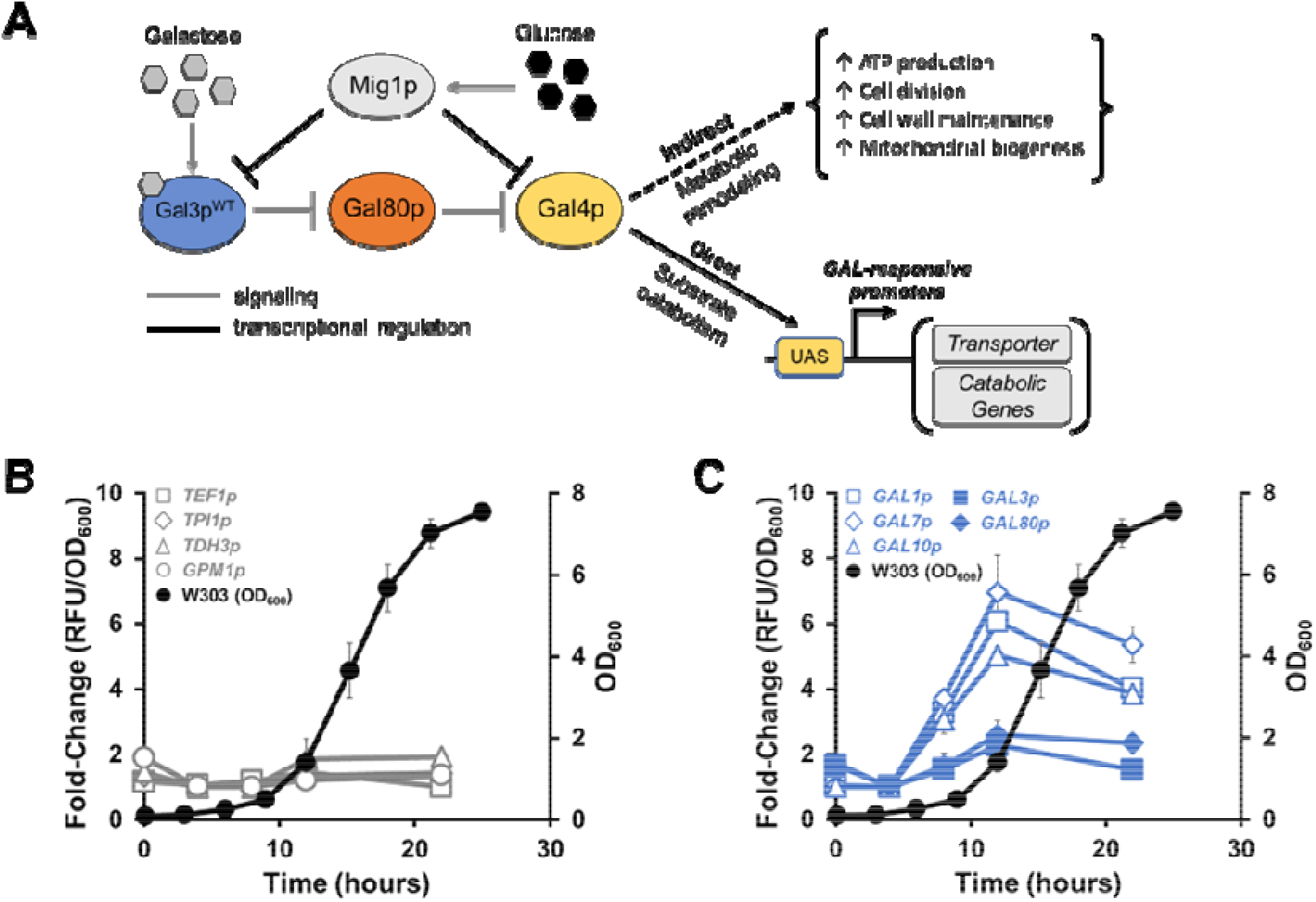
Expression from GAL-inducible promoters is coordinated with growth. **(A)** Simplified regulatory overview of the GAL regulon. The binding of galactose to sensor protein Gal3p enables it to relieve the repression of Gal80p on Gal4p via protein-protein interactions. Gal4p binds to its cognate Upstream Activating Sequences (UAS) to directly transcribe galactose catabolic genes as well as to indirectly reshape global metabolism for rapid growth. The regulon is also repressed in the presence of excess glucose, a regulatory logic that ensures the regulon is activated only in the presence of its target carbon source but not when a preferred carbon source is present. Expression of EGFP in wild-type strain W303-1a from **(B)** strong constitutive promoters (*TEF1p*, *TPI1p*, *TDH3p*, *GPM1p*) and **(C)** GAL-inducible promoters (*GAL1p*, *GAL7p*, *GAL10p*, *GAL3p*, *GAL80p*) at different time points during cultivation on galactose overlaid with cell density (OD_600_). Each data point represents the average of three (for OD_600_ values) or four (for fluorescence) biological replicates ± sd.

In this study, we demonstrate that, in addition to the known genome-wide changes in gene expression that occur upon GAL regulon activation, the expression from GAL-inducible promoters is dynamic and appears to be coordinated with growth whereas expression from constitutive promoters is constant. Hypothesizing that both aspects of this metabolic coordination may be universally beneficial for the assimilation of non-native substrates, we sought to develop this semi-synthetic regulon system into a platform approach that enables the rapid engineering of efficient synthetic heterotrophy without having to continually re-engineer substrate-specific induction. To do so, we created a variant of sensor protein that we term Gal3p^MC^ (i.e., <u>m</u>etabolic <u>c</u>oordinator) and demonstrate that it, unlike the wild-type Gal3p (Gal3p^WT^), can strongly activate the GAL regulon on numerous structurally diverse substrates. We show that Gal3p^MC^ can recapitulate the dynamic activation of the native regulon and, through the use of molecular dynamics and metadynamics simulations, that the mutations it carries enable it to do so in an inducer-independent manner. We found that this substrate-agnostic system retains the benefits of substrate-specific activation without any undue burden when engineering synthetic heterotrophy. We also show that using Gal3p^MC^ to implement a REG approach to growth on three non-native substrates – xylose, arabinose, and cellobiose – yields superior performance when compared to a CONS approach. Finally, we provide a first demonstration that a single strain expressing Gal3p^MC^ is capable of rapid, simultaneous, and complete utilization of all three non-native substrates concurrent with rapid growth – paving the way for a universal synthetic heterotrophy platform.

## RESULTS

### Expression under the GAL regulon is dynamic and recapitulates growth phase

Existing efforts to engineer synthetic heterotrophy generally utilize strong, constitutive promoters to control the expression of the necessary catabolic genes. Constitutive promoters are natively able to recruit cellular transcriptional machinery to yield a roughly consistent expression level independent of cellular state and environmental context. This ‘always on’ phenotype contrasts with the control of catabolic gene expression observed in many regulatory structures that coordinate native substrate utilization in response to nutrient availability, including the GAL regulon of *S. cerevisiae*. To understand how these different approaches could impact the efficiency of substrate utilization, we used a fluorescent reporter (*EGFP*) to quantify the expression dynamics of several GAL-responsive (*GAL1p*, *GAL7p*, *GAL10p, GAL3p,* and *GAL80p*) and constitutive (*TPI1p*, *TEF1p*, *TDH3p*, and *GPM1p*) yeast promoters in a wild-type strain (W303-1a) that contains the native GAL regulon. During growth on galactose, we found that expression from the constitutive promoters was relatively stable across a 24-hour period (varying less than two-fold) (**Figur 1B**). In contrast, expression from GAL-responsive promoters starts lower (∼50% of constitutiv promoters) but increases by ∼4-6 fold for the promoters that control the expression of the galactose catabolic (Leloir pathway) genes (*GAL1p*, *GAL7p*, and *GAL10p*) while remaining relatively constant for the promoters controlling regulatory genes (*GAL3p* and *GAL80p*) (**Figure 1C**). When we overlaid the OD_600_ profile, we observed that expression from Leloir pathway promoters tracks with growth phase, with the lowest expression levels occurring when the cells are in lag phase (t ∼ 0 h), maximum expression during the exponential growth phase, and finally diminishing expression as galactose is depleted and the cells enter stationary phase (t > 20 h). During growth on glucose, the expression profile of the constitutive promoters was largely unchanged (**Figure S1A**) while expression from GAL-inducible promoters is strongly repressed because of carbon catabolite repression (**Figure S1B**). Overall, GAL-responsive promoters are more dynamic and stronger (upon full activation) relative to constitutive promoters ^21, 22^.

### Gal3p^MC^ activates the GAL regulon in the presence of structurally diverse carbon sources

While the creation of a xylose-inducible sensor protein variant Gal3p^Syn4.1^ enabled us to adapt the GAL regulon for growth on xylose ^18^, we wanted to develop a method for rapidly applying a regulon approach to other non-native substrates without having to repeatedly re-engineer substrate-specific sensing. Blank et al. previously reported various autoactivating Gal3p variants but we found that the best among those (*viz.* F237Y, S509P) could only partially activate GAL promoters without galactose (**Figure S2A**) ^23^. To enable complete activation, equivalent to the WT GAL regulon with galactose, we combined the F237Y mutation to our Gal3p^Syn4.1^ variant ^24^ to yield a mutant we term Gal3p^MC^ (<u>m</u>etabolic <u>c</u>oordinator) (**Figure S2B**). To compare the ability of Gal3p^WT^ and Gal3p^MC^ to activate the regulon in the presence of a given carbon source, we monitored the fluorescence resulting from a *GAL1p-yEGFP* reporter construct (**Figure 2A**) in the presence of numerous structurally distinct carbon sources substrates – galactose, arabinose, xylose, cellobiose, raffinose, ethanol/glycerol, sucrose, and glucose – in a strain (VEG16) that lacks the galactose sensing and catabolic genes (Δ*GAL3;* Δ*GAL1;* Δ*GAL10;* Δ*GAL7;* Δ*GRE3)* but retains *GAL80* and *GAL4*. We performed this experiment using two native carbon sources, raffinose (**Figure 2B**) or sucrose (**Figure S3**), to support cell growth. We observed that Gal3p^WT^ strongly activates the regulon in the presence of its native inducer galactose and to a lesser extent xylose and arabinose, likely due to the structural similarity between these substrates. The indistinguishable signal between Gal3p^WT^ and th Δ*GAL3* control in presence of glucose reflects carbon catabolite repression while the small increase in fluorescence on raffinose and cellobiose likely reflects the weak activation known to result from overexpression of Gal3p^WT^ in the absence of glucose ^25^.

**Figure 2:**
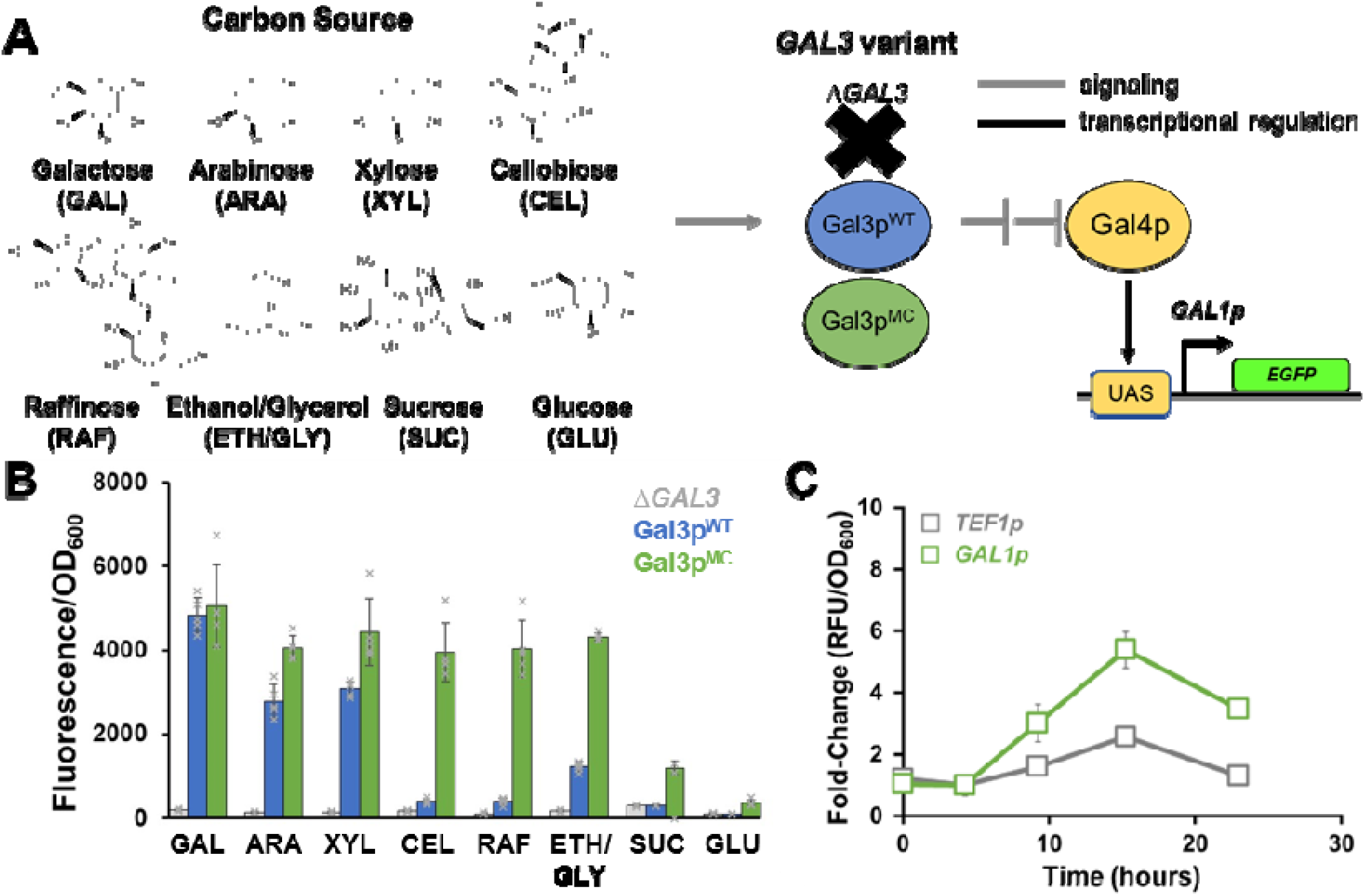
Sensor protein variant Gal3p^MC^ enables robust GAL regulon activation in an inducer-independent manner. **(A)** Schematic of fluorescent reporter assay used to quantify GAL regulon activation by different *GAL3* variants. Cells expressing a given variant (Δ*GAL3*, Gal3p^WT^, or Gal3p^MC^) are grown in the presence of both native and non-native substrates. Regulon activation drives EGFP expression from the GAL-inducible promoter, *GAL1p*. **(B)** Comparison of normalized fluorescence resulting from interaction between *GAL3* variant and carbon source (2% w/v) using RAF to support growth on substrates GAL, ARA, XYL, and CEL. SUC and GLU are repressing substrates. **(C)** Expression of EGFP from *TEF1p* and *GAL1p* during growth on galactose in a strain (SFS6) expressing Gal3p^MC^ to activate the regulon. Each data point represents the average of 5–6 biological replicates ± sd.

In contrast, Gal3p^MC^ activates the GAL regulon in the presence of all the non-native substrates to a similar extent as Gal3p^WT^ on galactose while remaining repressible by glucose and sucrose (**Figure 2B**), a result that suggests Gal3p^MC^ may be capable of recapitulating the dynamic catabolic gene expression profile and genome-wide expression changes known to support optimal growth on the native substrate galactose during growth on non-native substrates ^26^. To determine if regulon activation using Gal3p^MC^ remains dynamic, we constructed a strain (SFS6) that contains copies of *GAL3^MC^* integrated into the chromosome under the control of both *GAL1p* and *GAL3p*. This dual-feedback loop configuration mimics the organization of the native GAL regulon and has been demonstrated to increase both the overall magnitude and the homogeneity of activation to better effect a switch-like behavior among cells in the population ^18, 27^. As SFS6 lacks the Leloir pathway (Δ*GAL1/7/10*), these genes were supplied under the control of their native promoters on a plasmid (pRS423-GAL-REG) to facilitate growth on galactose. Using strain SFS6, we assessed the expression profile of the constitutive *TEF1p-EGFP* and GAL-responsive *GAL1p*-*EGFP* constructs during growth on galactose (**Figure 2C**). While expression from *TEF1p* was observed to increase by 2–3-fold at one time point, it generally stayed within a two-fold range whereas expression from *GAL1p* appeared to mimic the profile observed upon activation by the wild-type regulon (**Figure 1C**), exhibiting a >5-fold increase in fluorescence.

### Molecular simulations support inducer-independent regulon activation by Gal3p^MC^

Experiments have shown that the binding of Gal3p^WT^ to its ligand galactose induces and stabilizes a conformational change from an ‘open’ to a ‘closed’ state that facilitates interaction with Gal80p and subsequent de-repression of Gal4p ^15^. However, mapping the mutations carried by Gal3p^MC^ onto a structure of Gal3p^WT^ in complex with Gal80p (PDB: 3V2U) reveals that the mutations are distributed throughout the protein (**Figure S4**). Combined with the structural diversity of substrates (pentoses, hexose, disaccharide, and trisaccharide) on which Gal3p^MC^ exhibits robust regulon activation, observations suggest that Gal3p^MC^ may adopt a ‘closed’ conformation that enables it to bind Gal80p and relieve repression on Gal4p without requiring interaction with a ligand/inducer. As the transition between open and closed states is associated with the relative movement of two domains in Gal3p^WT^, while a third ‘hinge’ region remains relatively static, we decided to explore this inducer-independent activation hypothesis using molecular dynamics (MD) simulations. To do so, we monitored the distance between two residues (D104 and L370) found on opposing moving domains (**Figure S5A**) over the course of 1000 ns molecular dynamics (MD) simulations of Gal3p^WT^ in complex with ATP and Mg^2+^ without galactose (WT^Gal–^), Gal3p^WT^ in complex with ATP, Mg^2+^, and galactose (WT^Gal+^), and Gal3p^MC^ in complex ATP and Mg^2+^ without galactose (MC^Gal–^). Simulations revealed that distances between the residues were more similar between MC^Gal–^ and WT^Gal+^ (**Figure S5B**). Calculations of active site cavities at the end of the 1000 ns simulation also revealed a similar trend (**Figure S5C–E**). To understand which residues most contributed to the dynamics observed in each of the three systems during these simulations, we performed cross-correlational studies of the relative movement between the residues of WT^Gal–^, WT^Gal+^, and MC^Gal–^ (**Figure S5F–H**). The lower intensities of residue-residue correlation in WT^Gal+^ and MC^Gal–^ maps indicate relatively stable dynamics relative to those of WT^Gal–^. To visualize the domain movements in detail, we used principal component analysis (PCA) for all three systems across the 1000 ns MD trajectories. Porcupine plots generated based on the extreme projections of the PCA show the intensity and direction of the motion of the C atoms in each structure. WT^Gal–^ showed highly dynamic conformational states while MC^Gal–^ and WT^Gal+^ showed relatively limited dynamics across the structure (**Figure S5I–K**).

Initializing the molecular dynamics simulations of the three conditions in the ‘closed’ conformation introduces a bias that prevents each system from adopting a range of conformational states. Metadynamics-based simulations use a potential bias to move a system out of a local energy minimum enabling the calculation of the free energy surface (FES) across a range of conformations. To perform these complementary metadynamics analyses, we began to define the conformational landscape to be explored by identifying centers of mass (COM) for each of the three Gal3p domains (C1–C3). Next, we defined the two collective variables (CV) to be manipulated during the simulation as (i) the distance between the COM of the two moving domains (C1 and C2) and (ii) the angle formed by the three COM at the hinge region (C3) (**Figure 3A)**. Examining the resulting FES for each system reveals differences in the CV values that yield energetically favorable structural conformations. The FES of WT^Gal–^ (**Figure 3B**) showed two Gaussian wells, one for a short-lived partially open state and one for an exceptionally prominent minimum corresponding to an irreversibly collapsed closed conformation catalyzed by strong interaction networks between the two moving domains (**Figure S6A**) not observed between the moving domains in the WT^Gal+^ (**Figure S6B**) and MC^Gal–^ (**Figure S6C**) systems. Therefore, it can be called misfolded state (W_m_) of the protein as the recovery of an open conformation would be highly energy-consuming, and therefore, unlikely. The FES of WT^Gal+^ showed a path of conformational transitions that leads from a closed (G_c_), but not collapsed, state to a fully open state (G_o_) of the protein (**Figure 3C**). Significantly, the closed state of WT^Gal+^ (G_c_) is structurally distinct from the closed, collapsed/misfolded state (W_m_) adopted by WT^Gal–^ suggesting that the latter is unlikely to interact with Gal80p in a productive manner. As the metadynamics simulation progresses, WT^Gal+^ eventually adopts an open state (G_o_) that visualization studies reveal to be a consequence of substate egress. The FES of MC^Gal–^ (**Figure 3D**) showed a single predominant minimum corresponding to a closed state (M_c_) that closely resembles the closed state (G_c_) of WT^Gal+^. These results suggest that the mutant Gal3p^MC^ forms a stable closed conformation closely resembling the closed WT^Gal+^ conformation, with a similar interaction network observed between the moving domains. When combined with the previously observed robust activation on structurally diverse substrates, the similarity between the closed state conformations adopted by WT^Gal+^ and the MC^Gal–^ systems suggests that Gal3p^MC^ can bind and sequester Gal80p in an inducer-independent manner. Recognizing that this substrate-agnostic activation phenotype of Gal3p^MC^ represents a departure from the inducer-dependent regulatory logic of the native regulon, we decided to test the fitness consequences of this transition when utilizing a regulon approach to engineer synthetic heterotrophy.

**Figure 3:**
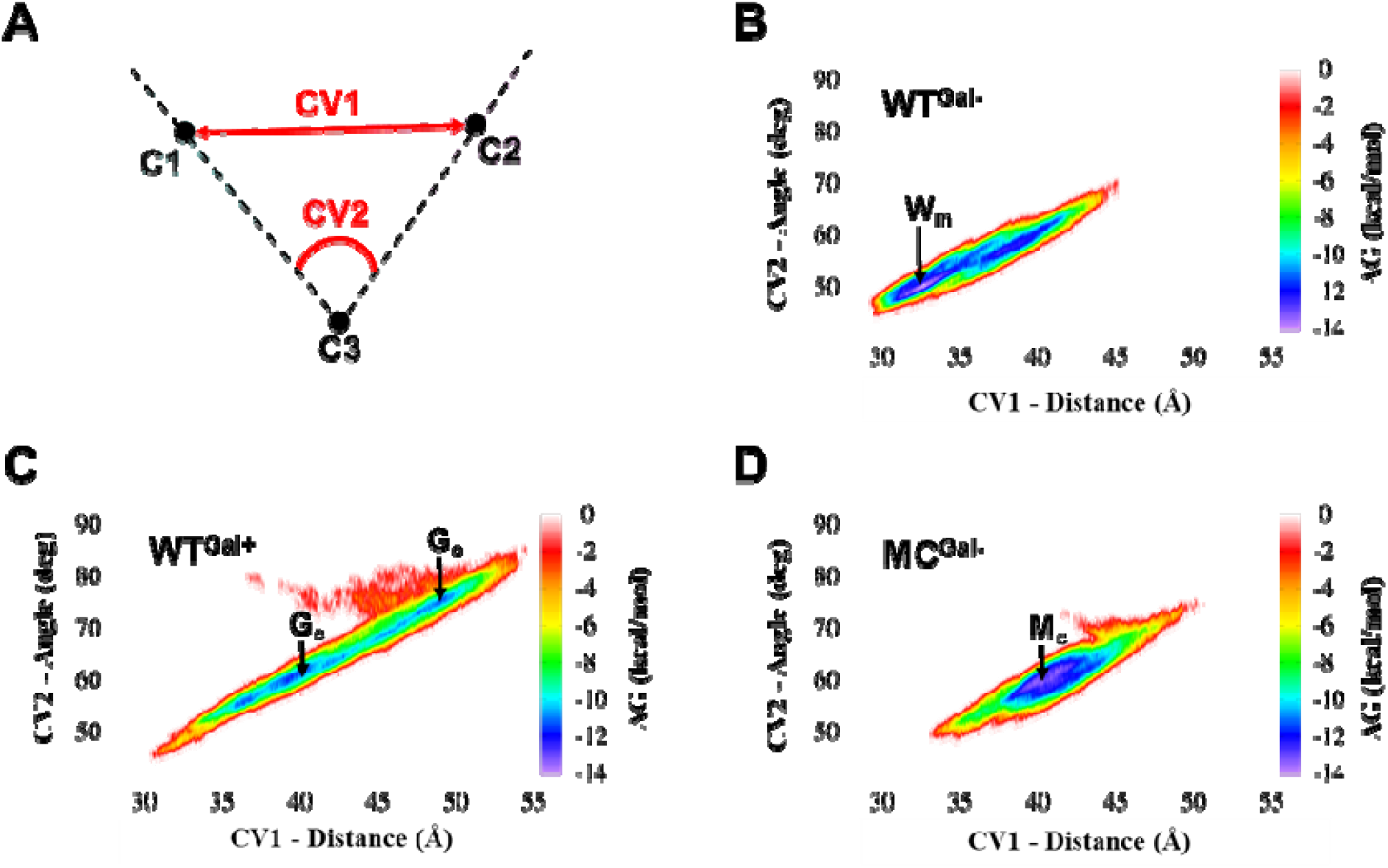
Metadynamics simulations identify energetically favorable conformations. **(A)** Two collective variables (CV) were defined to enable energetic exploration of the Gal3p conformational landscape: **CV1** as the distance between C1 and C2, the centers of mass of the two moving domains, and **CV2** as the angle formed between C1, C3 (the hinge region center of mass), and C2. Calculating the energy associated with different values of CV1 and CV2 enabled construction of the free energy surfaces (FES) for the **(B)** WT^Gal–^ (W), **(C)** WT^Gal+^ (G), and **(D)** MC^Gal–^ (M) systems. The subscripts “m”, “c”, and “o”, corresponding to misfolded (collapsed), closed, and open states, respectively, are used to associate prominent energetic minima with distinct conformations of Gal3p variants. Significantly, the most favorable conformations in the WT^Gal+^ and MC^Gal–^ systems correspond to a closed state that likely favors interaction with Gal80p and thus regulon activation.

### Interactions between Gal3p^MC^ with Gal80p are energetically favorable

We compared the energetics of Gal3p-Gal80p binding using the minimum energy conformations (W_m_, G_c_, M_c_) obtained from metadynamics-based simulations. Docking simulations between Gal3p and Gal80p were run for 50 ns and binding energy was calculated every 0.02 ns (**Figure 4A**). Among the complexe formed, the WT^Gal+^−Gal80p was energetically the most like the crystal structure complex (**Figure 4B**). The results also revealed that the complex formed by WT^Gal–^ with Gal80p was significantly less energetically favorable than the complexes formed by WT^Gal+^ and MC^Gal–^. This data further supports the idea that Gal3p^MC^ may be able to activate the GAL regulon in a substrate independent manner.

**Figure 4:**
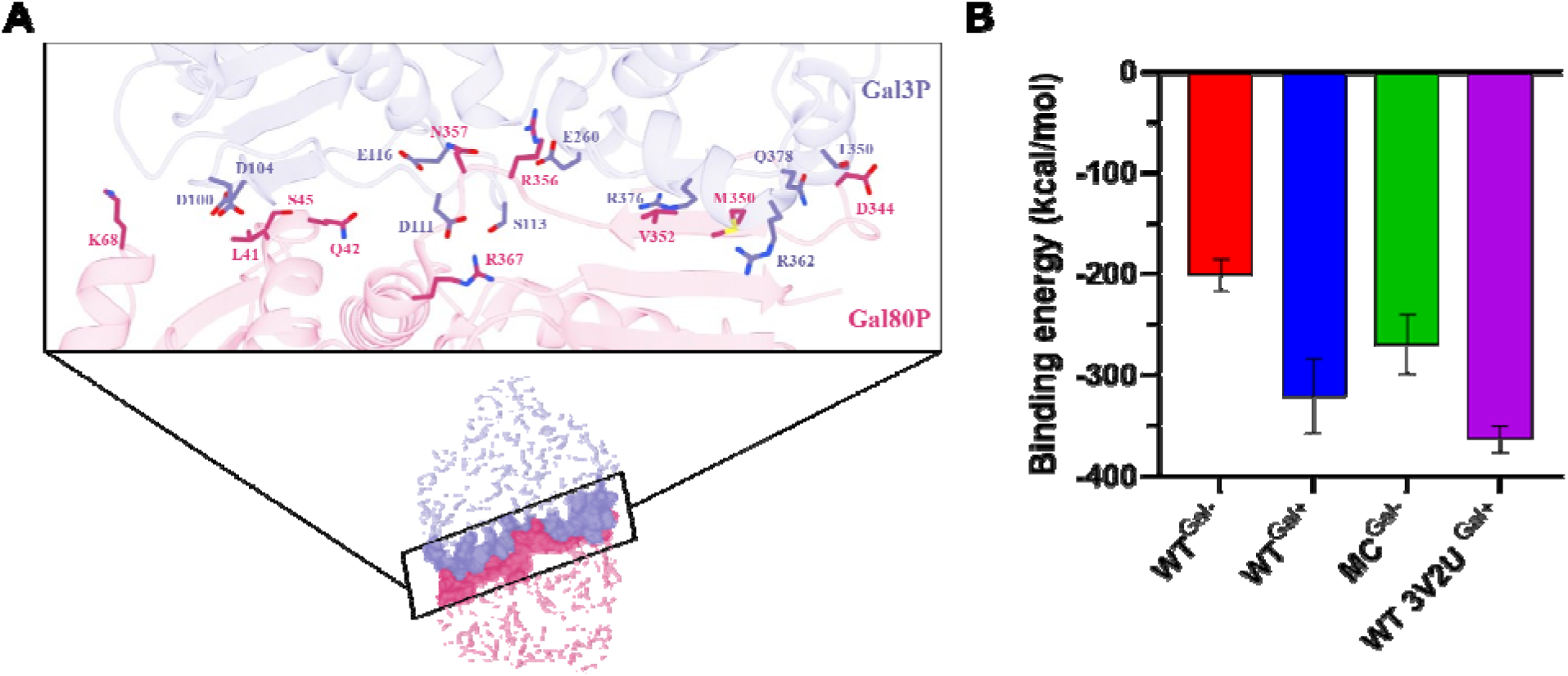
Docking interactions between Gal3p and Gal80p. **(A)** Detailed representation of the interface between Gal3p (purple) and Gal80p (pink) complex. The residues involved in the interface are highlighted in the call-out. **(B)** The average binding energy between Gal3p and Gal80p was calculated by sampling every 0.02 ns from the docked Gal3p-Gal80p complexes obtained through protein-protein docking study. A statistical analysis was performed on the average of the four binding energy values using ANOVA to determine whether there was a significant difference in the four sets of calculated binding energies. The results of the analysis showed an F-value of 2137 with 3 degrees of freedom and p-value of <0.0001, indicating a significant difference between all four.

### Substrate-agnostic and substrate-specific GAL regulon activation are equivalent

During our previous effort to adapt the GAL regulon to growth on xylose, regulon activation was accomplished using the xylose-inducible sensor-protein variant Gal3p^Syn4.1^ ^18^. To compare the performance of inducer-dependent (substrate-specific) and inducer-independent (substrate-agnostic) approaches for engineering synthetic heterotrophy, we transformed a plasmid (pRS426-XYL-REG) encoding the isomerase pathway genes for xylose utilization (*XYLA*3* ^28^*, XKS1*) under the control of GAL-responsive promoters into strains engineered for (i) xylose-inducible regulon activation using Gal3p^Syn4.1^ (strain VDT13), (ii) substrate-agnostic regulon activation using Gal3p^MC^ (strain SFS6), and (iii) substrate-agnostic activation via deletion of *GAL80*, the master repressor of the GAL regulon (strain SFS11), as disruption of this regulatory gene has been demonstrated to yield constitutive expression of galactose catabolic genes and enhance galactose utilization ^29, 30^ (**Figure 5A**). While regulon activation in both SFS6 and VDT13 requires an interaction between Gal80p and their respective Gal3p variant (Gal3p^MC^ and Gal3p^Syn4.1^), activation in SFS11 is totally de-regulated due to the absence of Gal80p. In addition, all three strains contain two accessory genes found to improve pentose utilization – the transporter *GAL2*^2^*^.1^*^31^ and the transaldolase *TAL1* – integrated into the chromosome under the control of GAL-responsive promoters. The growth of these three strains in defined (SC) media containing the native carbon sources sucrose (**Figure 5B**), glucose (**Figure 5C**), or raffinose (**Figure 5D**) was monitored to identify the potential fitness defects of each approach during strain handling. While all three approaches reached similar final cell concentrations on each of the three substrates, the Δ*GAL80* condition took roughly twice the time to do so compared with Gal3p^Syn4.1^ and Gal3p^MC^ (>30 h vs. ∼18 h) suggesting that de-regulation of the regulon results in reduced fitness relative to strains in which the native regulatory organization is maintained. As sucrose and glucose are both capable of exerting repression on the GAL regulon ^19^, subculturing all three strains from these two native substrates into complex (YP) media containing the non-native substrate xylose should therefore result in growth-coupled expression of xylose catabolic genes and rapid growth. Upon subculturing from sucrose, we observed that all three approaches enabled growth. However, the lower growth rate seen for the Δ*GAL80* strain compared with cells carrying *GAL3^Syn4.^*^1^ or *GAL3^MC^* (μ = 0.14 ± 0.001 h^−1^, 0.21 ± 0.008 h^−1^, and 0.21 ± 0.001 h^−1^, respectively) combined with the drastically lower final cell density supports maintenance of native regulatory structure over abolished control (**Figure 5E**). Subculturing from glucose as an overnight carbon source yielded similar results (**Figure S7**). Significantly, the equivalent performance of *GAL3^Syn^*^4^.^1^ and *GAL3^MC^* backgrounds on both native and non-native carbon sources indicates that inducer-independent activation enables the metabolic coordination of a regulon approach to be applied for engineering synthetic heterotrophy in a substrate-agnostic manner provided that the native regulatory organization is maintained.

**Figure 5:**
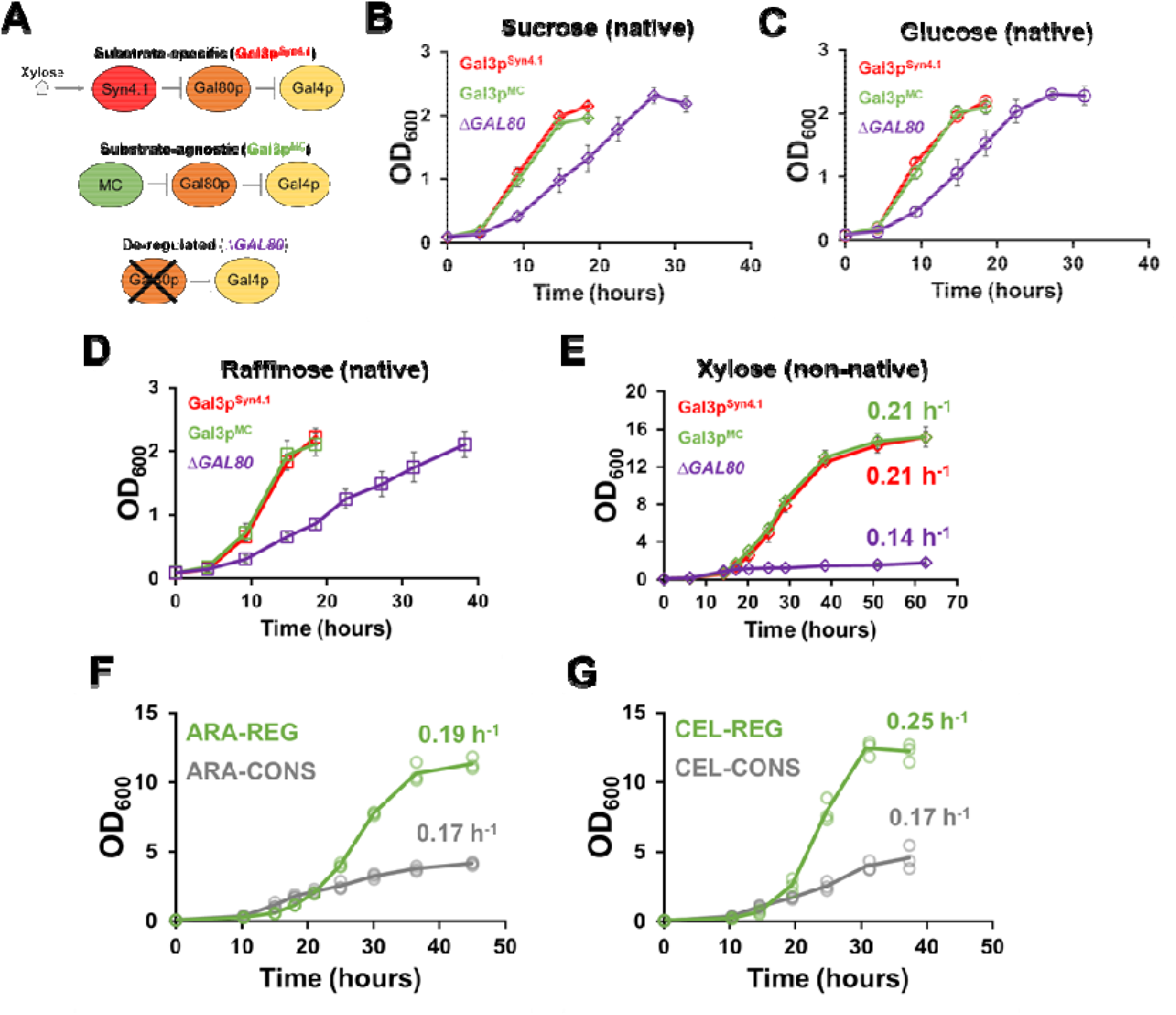
Comparing the fitness of substrate-specific and substrate-agnostic approaches to GAL regulon activation. **(A)** Schematic illustrating different ways of activating the GAL regulon in cells carrying xylose catabolic genes under the control of GAL-responsive promoters. Gal3p^Syn4.1^ requires induction by xylose, Gal3p^MC^ does not require an inducer but maintains the native regulatory architecture, while Δ*GAL80* effects inducer-independent activation by removing native regulatory elements. Growth of the three systems in defined (SC) medium containing 2% of **(B)** sucrose, **(C)** glucose, or **(D)** raffinose reveals that a de-regulated approach (Δ*GAL80*) reduces fitness on native carbon sources. **(E)** Comparing growth of the three GAL regulon activation systems on the non-native sugar xylose after being subcultured from sucrose. Comparison of constitutive (CONS) and regulon (REG) approaches to growth in complex media containing 2% of (**F**) arabinose or (**G**) cellobiose as sole carbon sources. All data points represent the average of three biological replicates ± sd.

To confirm that these results were not xylose-specific, we proceeded to use Gal3p^MC^ to compare REG and CONS approaches to growth on two additional non-native, lignocellulosic substrates – cellobiose and arabinose. To do so, we placed catabolic genes previously identified to enable the efficient utilization of arabinose (*araA*, *araB*, and *araD* from *Lactobacillus plantarum* ^32^) and cellobiose (*cdt-1* and *gh1-1* from *Neurospora crassa* ^33^) under the control of the GAL-responsive promoters *GAL1p*, *GAL10p*, and *GAL7p* and the strong, constitutive promoters *TEF1p*, *TPI1p*, and *GPM1p* to yield plasmids pRS426-ARA-REG, pRS426-ARA-CONS, pRS426-CEL-REG, and pRS426-CEL-CONS, respectively. Transforming pRS426-ARA-REG into *GAL3^MC^* integrant strain SFS6 (containing GAL-inducible copies of *GAL2^2.1^* and *TAL1*) yielded strain ARA-REG while transforming pRS426-ARA-CONS into strain VDT27 (that contains *GAL2^2.1^* and *TAL1* under the control of strong constitutive promoters) yielded strain ARA-CONS. Similarly, transforming pRS426-CEL-REG and pRS426-CEL-CONS into strains SFS3 and VEG16, respectively, that are comparable to SFS6 and VDT27 except that they lack accessory genes *GAL2^2.1^*and *TAL1*, yielded strains CEL-REG and CEL-CONS. The REG strain exhibited a higher growth rate than the CONS strain on both defined (0.16 ± 0.005 h^−1^ vs. 0.05 ± 0.005 h^−1^) (**Figure S8A**) and complex (0.19 ± 0.002 h^−1^ vs. 0.17 ± 0.003 h^−1^) (**Figure 5F**) media containing 2% arabinose as a sole carbon source. Similarly, the REG strain grew faster than its CONS counterpart on both defined (0.14 ± 0.001 h^−1^ vs. 0.09 ± 0.005 h^−1^) (**Figure S8B**) and complex (0.25 ± 0.002 h^−1^ vs. 0.17 ± 0.003 h^−1^) (**Figure 5G**) media containing 2% cellobiose as a sole carbon source. In general, we found that the strength of the promoters driving the catabolic gene expression corresponded well with growth phenotype (**Figure S9**). Taken together, these data demonstrate that the metabolic coordination effected by Gal3p^MC^ is broadly beneficial to growth on non-native carbon sources compared with the existing constitutive paradigm with minimal additional engineering.

### Gal3p^MC^ enables rapid and complete co-utilization of multiple non-native substrates

Having demonstrated that a regulon approach is beneficial to growth on all three non-native, lignocellulosic substrates individually, we sought to utilize the inducer-independent regulon activation of Gal3p^MC^ to test a REG approach to their co-utilization. As previous studies have observed that the division of catabolic labor among a consortium of strains can enable faster and/or more efficient co-utilization when compared with a single strain engineered to utilize all substrates simultaneously ^34–36^, we decided to test both approaches. To accommodate all three plasmids into a single strain, the catabolic genes for each substrate were moved to plasmids containing unique selectable markers to create pRS423-XYL-REG, pRS425-ARA-REG, and pRS424-CEL-REG. Transforming these plasmids into strains SFS6, SFS6, and SFS3, respectively, represented a three-strain (Consortium) approach, while transforming all three plasmids into SFS6 represented a single strain (Consolidated) approach (**Figure 6A**). While both Consortium and Consolidated approaches exhibited simultaneous co-utilization of all carbon sources and achieved similar final cell densities during growth on YP media containing 1% of each substrate, we were surprised to find that the Consolidated approach resulted in a faster maximum growth rate than the Consortium approach (0.20 ± 0.003 h^−1^ vs. 0.17 ± 0.003 h^−1^; two-tailed, two-sample t-test, t = 8.41, df = 3, p < 0.01) (**Figure 6B**). However, neither approach resulted in complete pentose consumption even after cellobiose was fully depleted. This led us to hypothesize that there may be flux imbalances in the pentose utilization pathways.

The success of combinatorial pathway design in optimizing both catabolic ^4,37, 38^ and anabolic pathways ^39, 40^ motivated us to test the performance of each of the six possible GAL-responsive promoter-catabolic gene pairings for arabinose (**Figure 6C**) and xylose (**Figure 6D**) in Gal3p^MC^ integrant strain SFS6. The arabinose constructs all performed similarly, with growth rates clustered in a small range of values (0.20 − 0.22 h^−1^). A similar range of growth rates was observed for the xylose constructs (0.20 − 0.21 h^−1^) but the substantially reduced final cell density of two of the arrangements (L-XK-XI and XK-L-XI) indicates that there can be fitness consequences to expression imbalances. As no clearly superior combination could be identified to the initial arabinose and xylose designs, we decided to re-test a Consolidated approach to co-utilization on 1% of each substrate (**Figure 6E**) using designs (ABD for arabinose, XI-L-XK for xylose) identified as optimal in a related investigation by our lab into efficient pentose assimilation ^19^. Hypothesizing that incomplete pentose utilization may also reflect limitations in the carrying capacity of the medium itself, we also tested different substrate ratios (cellobiose-xylose-arabinose: 0.7%–0.7%–0.7%, 1%–0.5%–0.5%, and 0.4%–0.8%–0.8%) (**Figure S10A-C**) at a lower (∼2%) total substrate concentration. Encouragingly, all substrates were utilized to undetectable levels (<0.05%, limit of detection) in approximately 50 h of cultivation for all conditions. For the 3% total sugar condition, we observed an 81% improvement in final cell density when using the optimized pentose plasmids compared with the non-optimized (OD_600_ = 17.2 vs. 9.5, respectively). For the three 2% total substrate conditions, the final cell density was 41% greater on average using the same comparison (OD_600_ ∼13.5 vs. 9.5, respectively). Despite their similar performance during growth on a single sugar, the higher final OD_600_ values and complete substrate utilization observed upon switching from the initial to the optimized designs suggests that the relative expression level of catabolic genes optimal for co-utilization may be different than for the single substrate. Considering that the highest single-sugar growth rates obtained in this study are either the best (for cellobiose) or competitive with the best (xylose, arabinose) aerobic growth rates observed in the literature (**Table S1**) without the need for extensive genome editing or adaptive laboratory evolution (ALE), we believe that together these results demonstrate the flexibility and effectiveness of utilizing a regulon approach to engineer efficient synthetic heterotrophy on diverse substrates.

**Figure 6:**
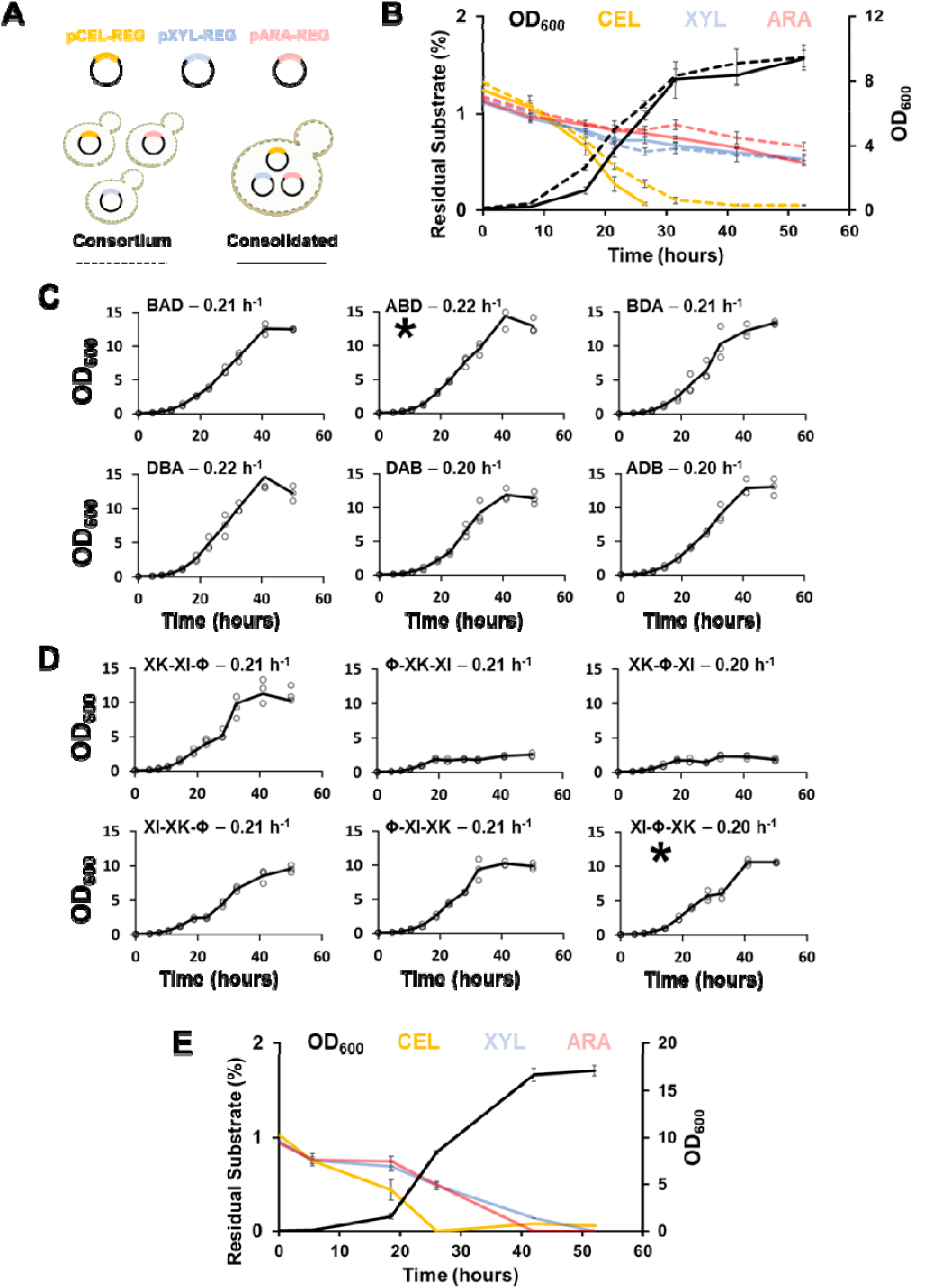
Gal3p^MC^ enables rapid and complete co-utilization of multiple non-native substrates. **(A)** Placing REG plasmids for the catabolism of cellobiose, xylose, and arabinose into separate strains represents a Consortium approach to co-utilization while placing all three plasmids into a single strain represents a Consolidated approach. **(B)** Cell density (black) and percent residual substrate of cellobiose (yellow), xylose (light blue), and arabinose (pink) of Consortium (dotted lines) and Consolidated (solid lines) approaches to co-utilization in complex media containing 1% of each substrate. Incomplete pentose utilization motivated the exploration of all six possible pairings between GAL-inducible promoters (*GAL1p*, *GAL10p*, and *GAL7p* in order) and the catabolic genes for **(C)** arabinose (A = L-arabinose isomerase, B = L-ribulokinase, D = L-ribulose-5-phosphate-4-epimerase) and **(D)** xylose utilization (XI = xylose isomerase, XKS = D-xylulokinase, L = no gene). Promoter-gene pairings used in subsequent co-utilization experiments are indicated by (**\***). **(E)** Re-testing a Consolidated approach to co-utilization using expression-balanced pentose plasmids results in complete utilization of all three substrates. Data points represent the average of three (panel **B**) or two (panel **E**) biological replicates ± sd.

## DISCUSSION

The existing paradigm for engineering synthetic heterotrophy consists of constitutive expression of catabolic genes that channel the carbon source into central carbon metabolism (CCM) and largely ignores cellular regulation. In contrast, the assimilation of native carbon sources is often controlled by regulatory structures called regulons that coordinate substrate catabolism with broader metabolism to optimize biomass formation. Activation of the galactose (GAL) regulon, for instance, results not only in the expression of the genes necessary for the catabolism of galactose but global metabolic remodeling to support efficient growth ^17, 20^. In previous work, we established that adapting the GAL regulon to growth on the non-native substrate xylose improved growth rate and substrate utilization when compared with a constitutive approach ^18^. This surprising result motivated us to explore the potential of a semi-synthetic regulon system as a general approach to engineering synthetic heterotrophy in the important industrial microbe *S. cerevisiae* so that it can be easily and rapidly applied to other attractive potential substrates for yeast biotechnology.

In this work, we demonstrate that expression from the GAL-responsive promoters that natively control the expression of galactose catabolic genes (Leloir pathway) is dynamic and mirrors growth phases whereas expression from constitutive promoters is constant throughout cultivation. Thus, the regulatory organization of the GAL regulon functions to avoid the burden of unproductive protein synthesis by only expressing the genes for galactose assimilation only when it is present. The conservation of cellular resources effected by this inducible phenotype represents a design principle that has also been learned by metabolic engineers. Early observations of growth inhibition resulting from the overproduction of heterologous protein ^41^ led to the use of separate culturing phases for growth and production and strategies for dynamically switching between them ^42^. This concept of dynamic flux control has subsequently been applied at finer resolution to biosynthetic pathways. Engineered circuits that feature a biosensor enable a cell to continually monitor some aspect of its internal state or external environment and automatically adjust their metabolism in response ^43^, resulting in improvements in titer, rate, and yield (TRY) ^44–47^. While the use of sugar-responsive biosensors has been proposed to engineer cells to respond to the carbon sources present in a given lignocellulosic hydrolysate for consolidated bioprocessing, the suggested implementations decouple sensing from the broader regulatory and metabolic context in which the sensors evolved ^48^. To our knowledge, our semi-synthetic regulon approach is the first time that this dynamic activation concept has been applied to engineering catabolism in a way that interfaces with native metabolism for the purpose of efficient biomass formation. This coordination is significant as the regulation of CCM has evolved to ensure that resource uptake and utilization are balanced ^3^, with the transcription level of a large fraction of the yeast genome (∼15 – 27%) ^49, 50^ known to be correlated with growth rate. We, therefore, posit that regulon control of catabolic gene expression, where the amount of carbon fed into CCM is innately connected to the utilization of those resources to match cellular needs, is likely beneficial to fast growth regardless of the carbon source being assimilated.

While we were previously able to engineer a sensor protein variant (Gal3p^Syn4.1^) capable of strong regulon activation upon induction with xylose, that effort involved an extensive directed evolution campaign that would have to be repeated for each new substrate ^18^. Here, we report the construction of a sensor protein variant, Gal3p^MC^ (<u>m</u>etabolic <u>c</u>oordinator), which robustly activates the GAL regulon on structurally diverse, non-native carbon sources to a similar degree as Gal3p^WT^ upon induction with galactose. Using metadynamics simulations, we show that the most energetically favorable conformation for Gal3p^MC^ without galactose (MC^Gal–^) is similar to the closed conformational state of Gal3p^WT^ in complex with galactose (WT^Gal+^). In contrast, the free energy surface of Gal3p^WT^ in the absence of galactose (WT^Gal–^) is characterized by a deep energy well reflective of the active site collapsing/misfolding, a conformation that is unproductive for signal transduction. Molecular dynamics (MD) simulations of these three systems reveal large-scale domain movements in the WT^Gal–^ system throughout the course of the simulation whereas the WT^Gal+^ and MC^Gal–^ systems exhibit much reduced and more localized movement indicating that the mutations carried by Gal3p^MC^ stabilize the protein in a closed state that enables interaction with Gal80p without requiring an inducer. The inducer-independent mechanism of Gal3p^MC^ would normally ablate the dynamic expression profile exhibited by the native, inducer-dependent regulon as it would constantly be turned on. However, we also observed repression of the Gal3p^MC^-controlled GAL regulon on repressing substrates (glucose, sucrose), which allows for dynamic activation. Gal3p^MC^ thus enables both the dynamic catabolic gene expression and global metabolic remodeling of the semi-synthetic regulon approach to be applied in a substrate-agnostic manner.

Although the inducer-independent activation effected by Gal3p^MC^ represents a departure from the inducer-dependent regulatory logic of the native GAL regulon that served as our inspiration, we, surprisingly, did not observe any major growth defects on either native or non-native substrates compared with a substrate-specific activation approach using Gal3p^Syn4.1^. In contrast, a Δ*GAL80* strain, which represents an alternative approach of achieving substrate-agnostic regulon activation, showed reduced fitness in both conditions, emphasizing the importance of maintaining the native regulatory organization when utilizing a regulon approach. During growth on the native carbon sources glucose, sucrose, and raffinose, the lower fitness likely results from the burden of constitutive GAL regulon activation.

As we previously observed with xylose ^18^, using Gal3p^MC^ to implement a regulon approach for growth on arabinose and cellobiose resulted in faster growth rates and higher final cell densities compared to the traditional constitutive overexpression approach despite expressing the same set of catabolic genes. Significantly, the growth rates achieved using this rational approach are comparable or superior to the highest aerobic growth rates for these substrates identified in the literature (**Table S1**) without the need for time-consuming and/or expensive experiments like ALE (adaptive laboratory evolution), functional genomics, and reverse genetics, or even deletion of genes previously found to mitigate stress of growth on pentoses (e.g., *PHO13, ALD6, ASK10, ADH6, COX4, CYC8*, etc. ^51^). The success of a semi-synthetic regulon approach in all the single-sugar conditions motivated us to explore their co-utilization as the simultaneous and complete co-utilization of the constituent sugars in lignocellulosic hydrolysates has been identified as essential to their valorization ^52^. One of the major challenges of engineering the co-utilization of lignocellulosic sugars is that the presence of glucose exerts carbon catabolite repression (CCR) in many organisms, hindering their ability to metabolize other sugars until glucose is depleted and repression is relieved ^53, 54^. However, Jin & Cate et al. found that CCR could be avoided by implementing a heterologous pathway for cellobiose, a glucose disaccharide that does not trigger CCR ^33, 55^. We demonstrate that using Gal3p^MC^ to implement a regulon approach enabled rapid and complete co-utilization of three non-native carbon sources – xylose, arabinose, and cellobiose – in a single strain in only ∼48 h. Despite being evolved for the catabolism of galactose, these results reinforce the universality and robustness of the metabolic coordination effected by the GAL regulon for engineering synthetic heterotrophy as well as the plasticity of yeast metabolism to diverse substrates. However, our findings do not discount the importance of existing metabolic engineering tools and strategies. For example, complete utilization of pentoses during co-utilization required balancing the expression levels of pathway genes. Additionally, while Gal3p^MC^ enables a regulon approach to be rapidly adapted to structurally diverse, non-native carbon sources, it is dependent on the development and importation of an efficient catabolic module. This requirement limits its immediate application to attractive renewable feedstocks like C1 compounds (i.e., CO_2_ ^56^, formate, methanol ^57^, methane ^58^) as efforts to engineer their assimilation in *S. cerevisiae* remain nascent. Researchers have noted the potential of platform strains that maximize the concentration of a key precursor to speed the construction and optimization of biosynthetic pathways ^59^. We envision that the integration of substrate catabolism with broader metabolism characteristic of a regulon approach could similarly accelerate the process of engineering catabolism by reducing the number of iterative, semi-rational genetic interventions currently needed to engineer efficient growth on a given non-native substrate. Together, these results establish the metabolic coordination of a regulon approach as a fast, flexible, and effective strategy for engineering synthetic heterotrophy and provide a tool (Gal3p^MC^) for adapting it to growth on structurally diverse non-native substrates.

## MATERIALS AND METHODS

### Strains and Plasmids

A full list of strains and plasmids used in this study can be found in **Table S2** and **Table S3**, respectively.

### Media and Transformation

Yeast strains were grown at 30 °C in either defined Synthetic Complete (SC) (Yeast nitrogen base (1.67 g/L), ammonium sulfate (5 g/L), complete supplement mixture without His, Leu, Ura and Trp (0.6 g/L) (Sunrise Science Products, Inc.) or complex Yeast Peptone (YP) medium (20 g/L Yeast Extract, 40 g/L Casein Peptone, 200 mg adenine) medium. SC media was augmented with appropriate nutrients to select for maintenance of plasmids carrying a given auxotrophy. Luria Bertani (LB) broth and LB agar plates with 100 μg/mL of ampicillin when required were used for all *E. coli* propagation and transformation experiments. Commercial *E. coli* strain NEB-5α (New England Biolabs) was used for MES transformation when constructing plasmids as well as for plasmid propagation. Upon isolation, plasmids were sequenced before being transformed into yeast strains using the protocol of Gietz ^60^ and recovered upon agar plates with the appropriate auxotrophy.

### Strain Construction

The yeast strain W303-1a was used for constructing all the strains used in the study. The generation of knockout strain VEG16 from W303-1a was described previously ^18^. All subsequent strains were derived from VEG16 via chromosomal integrations using a previously described set of disintegrator plasmids and associated protocols ^61^.

### Plasmid Construction

*S. cerevisiae* promoter, gene, and terminator sequences were amplified from the genome using Phusion Polymerase (ThermoFisher Scientific). *XYLA*3* DNA sequence was provided by Prof. Hal S. Alper (University of Texas at Austin). *GAL2*^2^.^1^ DNA sequence was provided by Prof. Bernard Hauer (University of Stuttgart, Stuttgart, Germany). Cellobiose utilization genes *cdt-1* and *gh1-1* were provided by Prof. Huimin Zhao (University of Illinois at Urbana-Champaign). Synthesis of DNA primers and Sanger sequencing of plasmid DNA was outsourced to Genewiz. Substrate utilization plasmids were created by combining promoter, catabolic gene, and terminator sequences into previously described yeast shuttle vectors containing either *CEN6* ^62^ or *2µ* ^63^ origins. All plasmids were constructed using either NEBuilder HiFi DNA Assembly or restriction enzyme digest and ligation using reagents purchased from New England Biolabs.

### Quantifying Promoter Dynamics via Fluorescence

Overnight inoculums were grown in 200 µL in 96-well plates containing the required dropout SC medium with glucose (2%) for 24 hours. Cultures were then subcultured (5 µL) into 200 µL SC medium containing the appropriate carbon source (2%) and returned to the incubator. Fluorescence (excitation at 488 nm and emission at 525 nm) and OD_600_ were measured at numerous time points in a Spectramax M3 spectrophotometer to obtain Fluorescence/OD_600_.

### Molecular Dynamics (MD) Simulations

Extended molecular dynamics and metadynamics simulation were conducted on three systems: Gal3p^WT^ complexed with ATP and Mg^2+^ (WT^Gal–^), Gal3p^WT^ complexed with galactose, ATP, and Mg^2+^ (WT^Gal+^), and Gal3p^MC^ complexed with ATP & Mg^2+^ (MC^Gal–^). To establish a common simulation starting point for the three systems, each of the protein structures was modeled into the closed state via homology modeling using Swiss Model ^64^ based on the experimentally determined closed state structure (PDB ID: 3V2U) as a template (**Figure S11**). Side chain conformations of the template were retained when generating models. Ramachandran plot was used to confirm that none of the side chains were in disallowed regions (**Figure S12**). A comparison of the crystal structure (PDB ID: 3V2U) with the homology model revealed low Root Mean Square Deviation (RMSD) of 0.09 (**Figure S13**). Molecular dynamic simulation of the three different experimental setups was performed for 1000 ns using GROMACS–2019.4 ^65^. The molecules were parameterized using antechamber package from AMBER ^66^. Quantum polarized charges were derived using the GAMESS package ^67^ with 6-31G basis set and B3LYP level of theory AMBER99SB ^66^ force field was used to perform a simulation with the TIP3p water model. To solvate the complex, the TIP3P water model with a triclinic box volume of 1000 nm^3^ was used to mimic or replace the natural environment around the protein. The system was neutralized by adding Na^+^ ions to the system, 16574 water molecules and ions overall. Energy minimization of the system was performed using the steepest descent algorithm with convergence energy cut-off of 1000 kJ mol^−1^ with 1000 steps. Equilibration of the system was performed using NVT and NPT, where NVT is for constant volume and temperature and NPT is for constant pressure. The NVT and NPT were carried out for 500 PS with 300 K temperature using a Berendsen thermostat and the system was prepared for Molecular dynamic simulation for 1 µs at 300 K temperature. The system was made to run for 1000 ns, for every 100 ns the trajectory was saved for analysis. The first 100 ns were excluded for every analysis. The distance between C_α_ atoms of L370 and D104 has been calculated across the simulation 1000 ns long trajectory for all three proteins to know the variation in the movement of two domain regions across the time. RMSD (Root mean square Deviation) and RMSF (Root mean square fluctuation) of the backbone atoms were calculated over time. Using WORDOM tools ^68^, cross-correlation studies of residue-residue displacements along the trajectory were calculated for each system to quantify dynamic changes of across the moving domains. Similarly, principal component analysis (PCA) was calculated for each 100-1000 ns trajectory and porcupine plots were generated to analyze the intensity and movement of C_α_ atom. Visualization of structures was done using UCSF Chimera ^69^ and Visual Molecular Dynamics (VMD) ^70^.

### Metadynamics Simulations

Metadynamics-based simulations use a potential bias to move the system out of a local minimum, which is then used to explore the resulting free energy landscape (FES). Metadynamics-based simulations were conducted using the PLUMED software package ^71^ and GROMACS toolkit for all three experimental setups (WT^Gal-^, WT^Gal+^, and MC^Gal-^) after thorough equilibration obtained from NVT, NPT ensembles. C1, C2, and C3 are Centers of Mass (COMs) determined for the three regions of Gal3p that fall within the residues regions as shown in **Figure S5A**. One of the Collective Variables (CV) for metadynamics simulation was defined as the distance between C1 and C2. The other CV was defined as the angle between the intersection point of the vectors that pass through C1, C3 and C2, C3, respectively. Different simulations in this study were performed for 50 ns with a bias factor set to 8.0, the temperature set to 300 K, and the results were calculated accordingly. The free energy landscape was explored as the distance between the two domains and the change in the angle formed by the three COMs were varied. A superposition of modeled closed form and apo form was used to visualize the structural changes (**Figure S14**).

### Gal3p-Gal80p Docking Studies

Crystal structure of Gal80p complexed with the closed Gal3p structure (PDB ID 3V2U) was used as the reference for protein-protein docking studies. Protein-protein docking studies were conducted using PatchDock ^72, 73^ and then refined using Fiberdock ^74, 75^. The minimum energy conformations of WT^Gal+^, WT^Gal–^, and MC^Gal–^ attained from the metadynamics simulations were docked with Gal80p and the complexes obtained from protein-protein docking studies were simulated using GROMACS–2019.4 ^76^ for 50 ns ×2 to better equilibrate the complexes that were obtained. The coordinates of the complexes were sampled every 0.02 ns and the binding affinity in kcal/mol between the Gal3p and Gal80p structures for all three complexes was calculated using the molecular mechanics/Poisson-Boltzmann surface area (MM/PBSA) method ^77, 78^.

### Growth Studies

Overnight inoculums were grown in the required dropout SC medium with 2% sucrose (unless explicitly stated to be glucose/raffinose) for 24 h. The culture was washed twice in the growth medium (either SC or YP containing 2% of the target sugar) and resuspended at an initial OD_600_ of 0.1 in 250 mL shake flasks containing 20 mL of media. Cell growth OD_600_ measurement was checked at frequent time intervals (3–6 h) on SpectraMax M3 spectrophotometer (Molecular Devices). Growth rates (μ) for each biological replicate were quantified in Microsoft Excel using an exponential curve equation fit to at least three time points during logarithmic growth phase. These values were used to calculate the average growth rate and the standard error for a given condition.

### Residual Substrate Quantification

Residual substrate during co-utilization studies was measured using an Agilent HPLC system equipped with a Hi-Plex H-column and detected using 1260 Agilent ELSD detector. Mobile phase was 0.1% trifluoroacetic acid (TFA) with a flow rate of 0.6 mL/min. The ELSD detector’s nebulizer and evaporation temperature were set at 30 °C and nitrogen flow rate at 1.6 SLM (standard liter per minute). Using these settings, we observed characteristic retention times for cellobiose (9.3 min), arabinose (12.6 min), and xylose (11.8 min).

## DATA AVAILABILITY

All plasmids and strains used in this study can be obtained from the corresponding author under a material transfer agreement. Source data are provided with this paper.

## ACKNOWLEDGEMENTS

The authors would like to thank current and former Nair lab members for helpful discussions. This work was supported by NIH grant #DP2HD91798, #R21HD105934, and NSF grant #1935354, and Tufts Launchpad | Accelerator to N.U.N.

## AUTHOR CONTRIBUTIONS

N.U.N., S.F.S., V.D.T., and V.E.G. conceived and designed the research project. S.F.S., A.S., and N.U.N. co-wrote the manuscript. S.F.S., A.S., V.D.T., T.C.C., V.E.G., D.M.S., and T.B. performed the experiments. S.F.S., A.S., T.B., P.K.R., and N.U.N. analyzed the data. All the authors have reviewed the manuscript and approved it for submission.

