## Supplemental Information for "Towards universal synthetic heterotrophy using a metabolic coordinator"

**Table S1:** Aerobic growth rates of previously engineered *S. cerevisiae* strains on xylose, arabinose, and cellobiose including relevant genotypes.

| Substrate | Strain | Relevant Genotype | *μ* (h^-1^) | Reference |
| --- | --- | --- | --- | --- |
| Xylose | X389 | *PiXI, PsXK, HXT7m, PsTAL1, TKL1, RPE1, RKI1 ∆GRE3, ∆ASK10, ∆YPR1* | 0.26 | ^1^ |
|  | LI110A | *SsXR, SsXDH*, SsXK, SsRPE, SsRKI, SsTKL, SsTAL; Evolved* | 0.25 | ^2^ |
|  | XYL-REG (Syn4.1) | *xylA*3, XKS1, TAL1, GAL2^2.1^, GAL3^Syn4.1^ under GAL promoters, ∆GAL1/3/7/10, ∆GRE3* | 0.24 | ^3^ |
|  | YRH1114 | *PrXK, PrXI, codons optimized for S. cerevisiae; Evolved* | 0.23 | ^4^ |
|  | RWB 217 | *XYLA, XKS1, TAL1, TKL1, RPE1, RKI1, ΔGRE3; Evolved* | 0.22 | ^5^ |
|  | XYL-REG (MC) | *xylA*3, XKS1, TAL1, GAL2^2.1^, GAL3^MC^ under GAL promoters, ∆GAL1/3/7/10, ∆GRE3* | 0.21 | This Study |
|  | H131-A3^SB-1^ | *XYLA, PsXYL3, PsTAL1, TKL1, RPE1, RKI1; Evolved* | 0.197 | ^6^ |
|  | CMB.GS010 | *PsXYL1, PsXYL2, PsXYL3; Evolved* | 0.18 | ^7^ |
|  | SyBE005 | *XYL1, mXYL2, XKS1, RPE1, TAL1, RKI1, TKL1, ∆AUR1; Evolved* | 0.165 | ^8^ |
|  | UUU | *UbXR1, UbXDH, UbXK* | 0.15 | ^9^ |
|  | ATCC-F2 | *pgXR, pcXDH, aoXKS; Evolved* | 0.15 | ^10^ |
|  | ADAP8 | *XYLA, XKS, SUT1; Evolved* | 0.133 | ^11^ |
|  | SXA-R2P-E | *Ss_xyla*3, XKS1, Ss_tal1, ΔGRE3, ΔPHO13; Evolved* | 0.128 | ^12^ |
|  | YSX3-TAL1M | *XYL1, XYL2, PsXYL3, psTAL1* | 0.117 | ^13^ |
| Arabinose | ARA-REG (Syn4.1) | *Lp_araA, Lp_araB, Lp_araD, TAL1, GAL2-2.1, GAL3^Syn4.^ under GAL promoters, ∆GAL1/3/7/10, ∆GRE3* | 0.27 | ^3^ |
|  | IMX728 | *araA, araB, araD, Pc_araT, ∆GLK1, ∆HXK1, ∆HXK2, ∆GAL1, ∆GAL80, ∆GRE3, RPE1, TKL1, TAL1, NQM1, RKI1, TKL2* | 0.26 | ^14^ |
|  | ARA-REG (MC) | *Lp_araA, Lp_araB, Lp_araD, TAL1, GAL2-2.1, GAL3^MC^ under GAL promoters, ∆GAL1/3/7/10, ∆GRE3* | 0.22 | This Study |
|  | IMS0002 | *Piromyces xylA, Lp_araA, Lp_araB, Lp_araD, XKS1, TAL1, TKL1, RPE1, RKI1, ∆GRE3* | 0.15 | ^15^ |
|  | BWY1-SI | *araA (codon optimized), araBG361A, araD, GAL2; Evolved* | 0.112 | ^16^ |
|  | TMB3664 | *XKS1, TAL1, TKL1, RKI1, RPE1, XYL1^K270R^, XYL2, sLAD1(synthetic), sALX1(synthetic), ΔGRE3* | 0.05 | ^17^ |
|  | TMB3076 | *Bs_araA, Ec_araD, Ec_araB, XKS1, TAL1, TKL1, RKI1, RPE1, ΔGRE3* | 0.03 | ^18^ |
| Cellobiose | CEL-REG (MC) | *Nc_gh1-1, Nc_cdt-1, GAL3^MC^ under GAL promoters, ∆GAL1/3/7/10, ∆GRE3* | 0.25 | This Study |
|  | Y294[SFI] | *SfBGL1 (secreted using T. reesei xyn2 signal), ∆FUR1* | 0.23 | ^19^ |
|  | YE-Aa | *AaBGL1 (secreted using T. reesei xyn2 signal)* | 0.2005 | ^20^ |
|  | IBP1 | *Bs_bglP, SfBGL1* | 0.11 | ^21^ |
|  | YPH499 | *Nc_gh1-1, Nc_cdt-1* | 0.0341 | ^22^ |

**Table S2:** List of strains used.

| **Strain** | **Description** |
| --- | --- |
| W303-1a | *MATa leu2-3,112 trp1-1 can1-100 ura3-1 ade2-1 his3-11,15* |
| VEG16 | W303-1a Δ*GAL3;* Δ*GRE3;* Δ*GAL1;* Δ*GAL7;* Δ*GAL10* |
| VDT13 | VEG16 *lys2::GAL3p-GAL3^Syn4.1^-TEFt/ADH1t-GAL2-2.1-GAL1/10p-TAL1-HXT7t; leu2::GAL1p-GAL3^Syn4.1^-TEFt* |
| VDT27 | VEG16 *lys2::ADH1t-GAL2^2.1^-TEF1/TPI1p-TAL1-HXT7t* |
| SFS3 | VEG16 *lys2::GAL3p-GAL3^MC^-TEFt; leu2::GAL1p-GAL3^MC^-TEFt* |
| SFS6 | VEG16 *lys2::GAL3p-GAL3^MC^-TEF1t; leu2::GAL1p-GAL3^MC^-TEFt; his3::HXT7t-TAL1-GAL1/10p-GAL2^2.1^-ADH1t* |
| SFS11 | VEG16 *∆GAL80; lys2::HXT7t-TAL1-GAL1/10p-GAL2^2.1^-ADH1t* |
| XYL-REG (MC) | SFS6 *+* pVDT62 |
| ARA-REG (MC) | SFS6 + pVDT38 |
| ARA-CONS | VDT27 + pSFS3 |
| CEL-REG (MC) | SFS3 + pSFS22 |
| CEL-CONS | VEG16 + pSFS10 |

**Table S3:** List of plasmids used.

| **Appearance in Manuscript** | **Plasmid** | **Description** |
| --- | --- | --- |
| Expression dynamics - *GAL1p* | pVEG18-GAL1p | pRS426*, 2μ ori, URA3, GAL1p-EGFP-ADH1t* |
| Expression dynamics - *GAL7p* | pVEG20-GAL7p | pRS426*, 2μ ori, URA3, GAL7p-EGFP-ADH1t* |
| Expression dynamics - *GAL10p* | pVEG21-GAL10p | pRS426*, 2μ ori, URA3, GAL10p-EGFP-ADH1t* |
| Expression dynamics – *GAL3p* | pVEG19-GAL3p | pRS426*, 2μ ori, URA3, GAL3p-EGFP-ADH1t* |
| Expression dynamics – *GAL80p* | pVEG22-GAL80p | pRS426*, 2μ ori, URA3, GAL80p-EGFP-ADH1t* |
| Expression dynamics - *TEF1p* | pVEG24-TEF1p | pRS426*, 2μ ori, URA3, TEF1p-EGFP-ADH1t* |
| Expression dynamics - *TPI1p* | pVEG25-TPI1p | pRS426*, 2μ ori, URA3, TPI1p-EGFP-ADH1t* |
| Expression dynamics – *TDH3p* | pVEG27 | pRS426*, 2μ ori, URA3, TDH3p-EGFP-ADH1t* |
| Expression dynamics - *GPM1p* | pVEG28 | pRS426*, 2μ ori, URA3, GPM1p-EGFP-ADH1t* |
| Activation data - *∆GAL3* control | pVEG7 | pRS426*, 2μ ori, URA3, ADH1t-EGFP-GAL1p/GAL10p-KANMX-HXT7t* |
| Activation data - Gal3p^WT^ | pVEG8-WT | pVEG7*, 2μ ori, URA3,GAL3p-GAL3^WT^-TEF1t* |
| Activation data - Gal3p^MC^ | pVEG8-MC | pVEG7*, 2μ ori, URA3,GAL3p-GAL3^MC^-TEF1t* |
| Creation of MC integrant strains | pVDT43 | pIS385, *URA3, GAL3p-Gal3p-5.1-Teft* |
| Creation of MC integrant strains | pVDT44 | pIS376, *URA3, GAL1p-Gal3p-5.1-Teft* |
| Creation of pentose integrant strains | pSFS31 | pIS374, *URA3, HXT7t-TAL1-GAL1/10p-GAL2-2.1-ADH1t* |
| pRS426-XYL-REG | pVEG11 | pRS426*, 2μ ori, URA3, ADH1t-Piromyces_XYLA*3-GAL1p/GAL10p-XKS1-HXT7t* |
| pRS426-ARA-CONS | pSFS3 | pRS426*, 2μ ori, URA3, ADH1t-araB-TPI1/TEF1p-araA-Hxt7t-GPM1p-araD-TEF1t* |
| pRS426-ARA-REG | pVDT23 | pRS426*, 2μ ori, URA3, ADH1t-araB-GAL1/GAL10p-araA-HXT7t-GAL7p-araD-TEF1t* |
| pRS426-CEL-CONS | pSFS10 | pRS426*, 2μ ori, URA3, ADH1t-gh1-1-TPI1p/TEF1p-cdt-1-HXT7t* |
| pRS426-CEL-REG | pSFS11 | pRS426*, 2μ ori, URA3, ADH1t-gh1-1-GAL1p/GAL10p-cdt-1-HXT7t* |
| pRS423-XYL-REG | pSFS23 | pRS423*, 2µ ori, HIS3, ADH1t-XKS1-GAL1/GAL10p-XYLA-HXT7t* |
| pRS424-CEL-REG | pSFS22 | pRS424*, 2μ ori, TRP1, ADH1t-gh1-1-GAL1/10p-cdt-1-HXT7t* |
| pRS425-ARA-REG | pSFS18 | pRS425*, 2μ ori, LEU2, TEF1t-araD-GAL7p-HXT7t-araA-GAL1/GAL10p-araB-ADH1t* |
| BAD | pVDT14 | pRS416*, CEN ori, URA3, ADH1t- araB- GAL1p/GAL10p -araA-HXT7t-GAL7p-araD-TEFt* |
| ABD | pVDT38 | pRS416*, CEN ori, URA3, ADH1t- araA- GAL1p/GAL10p -araB-HXT7t-GAL7p-araD-TEFt* |
| BDA | pVDT39 | pRS416*, CEN ori, URA3, ADH1t- araB- GAL1p/GAL10p -araD-HXT7t-GAL7p-araA-TEFt* |
| DBA | pVDT40 | pRS416*, CEN ori, URA3, ADH1t- araD- GAL1p/GAL10p -araB-HXT7t-GAL7p-araA-TEFt* |
| DAB | pVDT41 | pRS416*, CEN ori, URA3, ADH1t- araD- GAL1p/GAL10p -araA-HXT7t-GAL7p-araB-TEFt* |
| ADB | pVDT42 | pRS416*, CEN ori, URA3, ADH1t- araA- GAL1p/GAL10p -araD-HXT7t-GAL7p-araB-TEFt* |
| ϕ-XK-XI | pVDT57 | pRS423*, 2µ ori, HIS3, ADH1t- XKS1- GAL10p/GAL1p -TEF1t -GAL7p- Piromyces_XYLA*3-HXT7t* |
| XK- ϕ-XI | pVDT58 | pRS423*, 2µ ori, HIS3, ADH1t- XKS1- GAL1p/GAL10p -TEF1t -GAL7p- Piromyces_XYLA*3-HXT7t* |
| XK-XI-ϕ | pVDT59 | pRS423*, 2µ ori, HIS3, ADH1t- XKS1- GAL1p/GAL10p -Piromyces_XYLA*3-HXT7t* |
| XI-XK-ϕ | pVDT60 | pRS423*, 2µ ori, HIS3, ADH1t- XKS1- GAL10p/GAL1p -Piromyces_XYLA*3-HXT7t* |
| ϕ-XI-XK | pVDT61 | pRS423*, 2µ ori, HIS3, ADH1t- XKS1- GAL7p-TEF1t -GAL1p/GAL10p - Piromyces_XYLA*3-HXT7t* |
| XI-ϕ-XK | pVDT62 | pRS423*, 2µ ori, HIS3, ADH1t- XKS1- GAL7p-TEF1t -GAL10p/GAL1p - Piromyces_XYLA*3-HXT7t* |

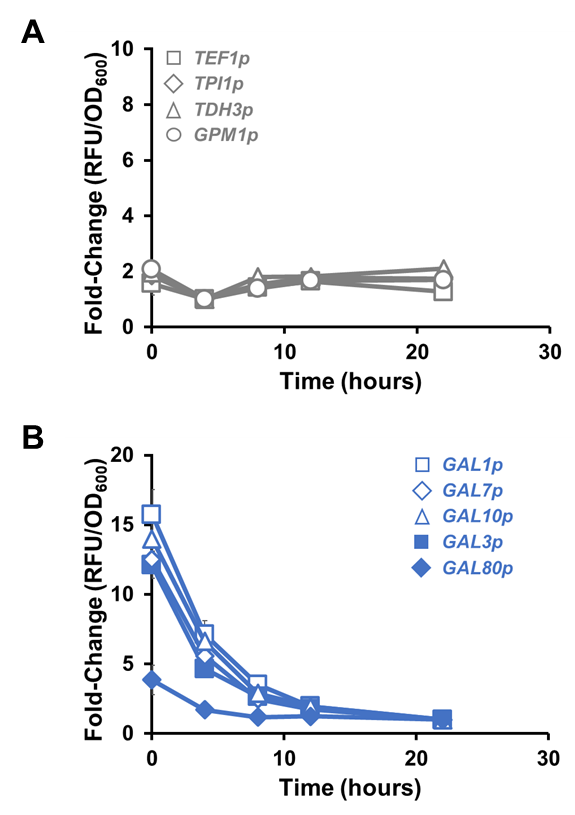

**Figure S1:** **Expression dynamics of constitutive and GAL-inducible promoters during growth on glucose.** Expression of EGFP in wild-type strain W303-1a from **(A)** strong constitutive promoters (*TEF1p*, *TPI1p*, *TDH3p*, *GPM1p*) and **(B)** GAL-inducible promoters (*GAL1p*, *GAL7p*, *GAL10p*, *GAL3p*, *GAL80p*) at different time points during cultivation on glucose. While expression profile from constitutive promoters is similar to that observed during growth on galactose, GAL-inducible promoters are repressed via carbon catabolite repression. Each data point represents the average of four biological replicates ± sd.

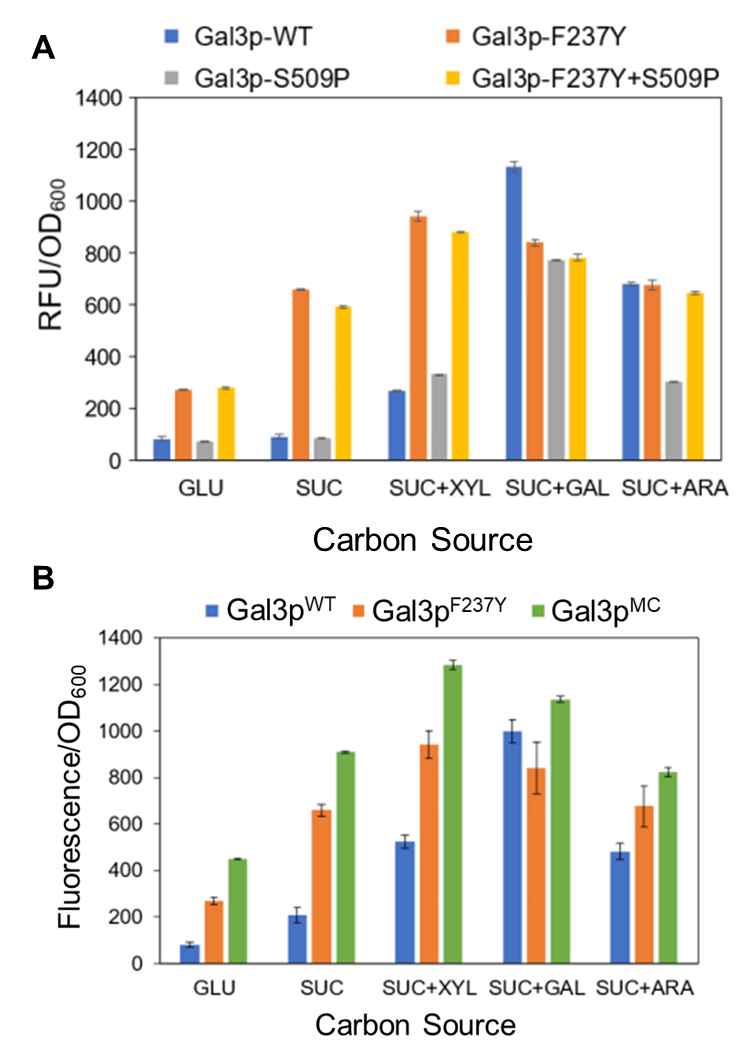

**Figure S2: Initial characterization of autoactivating Gal3p mutants. (A)** Activation of *GAL1p-EGFP* using partial autoactivating mutants identified by Blank et al. (1997). None of the 3 variants tested fully activate *GAL1p* to the same level as Gal3p^WT^ does with galactose. **(B)** Combination of F237Y with Gal3p^Syn4.1^ yielded Gal3p^MC^ that activates more strongly than F237Y alone. All experiments use 2% (w/v) sugar. GLU = Glucose, SUC = sucrose, GAL = galactose, ARA = arabinose, XYL = xylose. Each data point represents the average of three biological replicates ± sd.

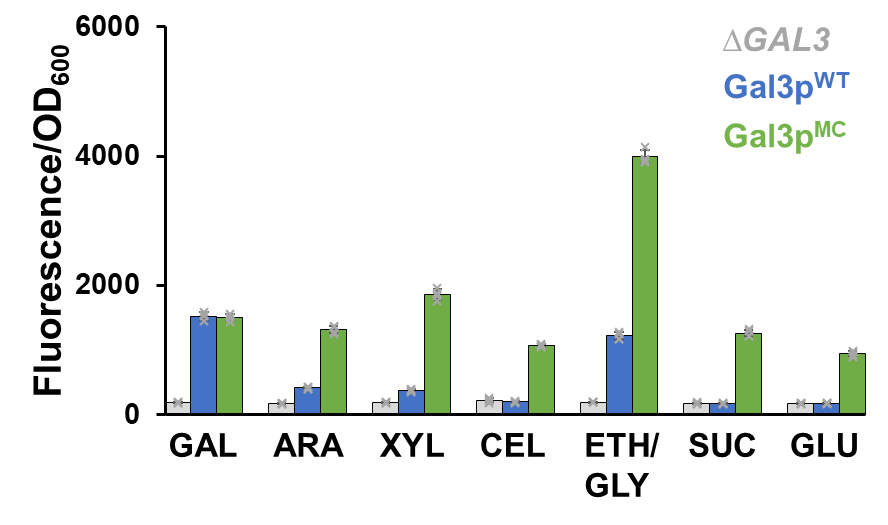

**Figure S3**: Comparison of fluorescence (normalized for cell density) resulting from interaction between *GAL3* variant and carbon source (2% w/v) using sucrose to support growth on substrates (GAL/ARA/XYL/CEL) for which strain VEG16 lacks catabolic genes. Each data point represents the average of four biological replicates ± sd. GAL = galactose, ARA = arabinose, XYL = xylose, CEL = cellobiose, ETH/GLY = ethanol/glycerol, SUC = sucrose, GLU = glucose.

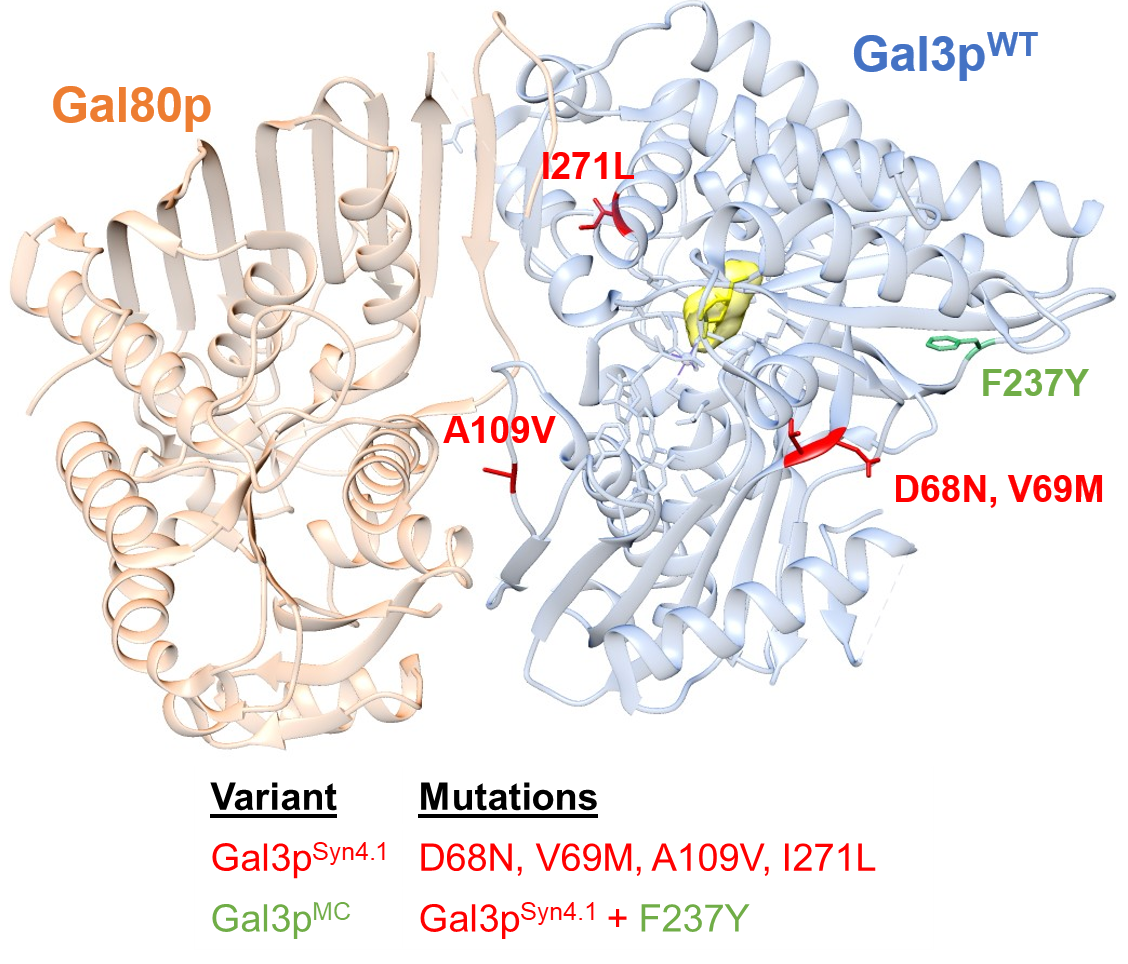

**Figure S4**: **Structure of wild-type Gal3p in complex with Gal80p.** Gal3p^MC^ was constructed by combining mutations identified as conferring robust xylose induction (D68N, V69M, A109V, I271L, highlighted in green) ^23^ in variant Gal3p^Syn4.1^ with a mutation (F237Y, highlighted in red) that confers a partial autoactivating phenotype ^24^.

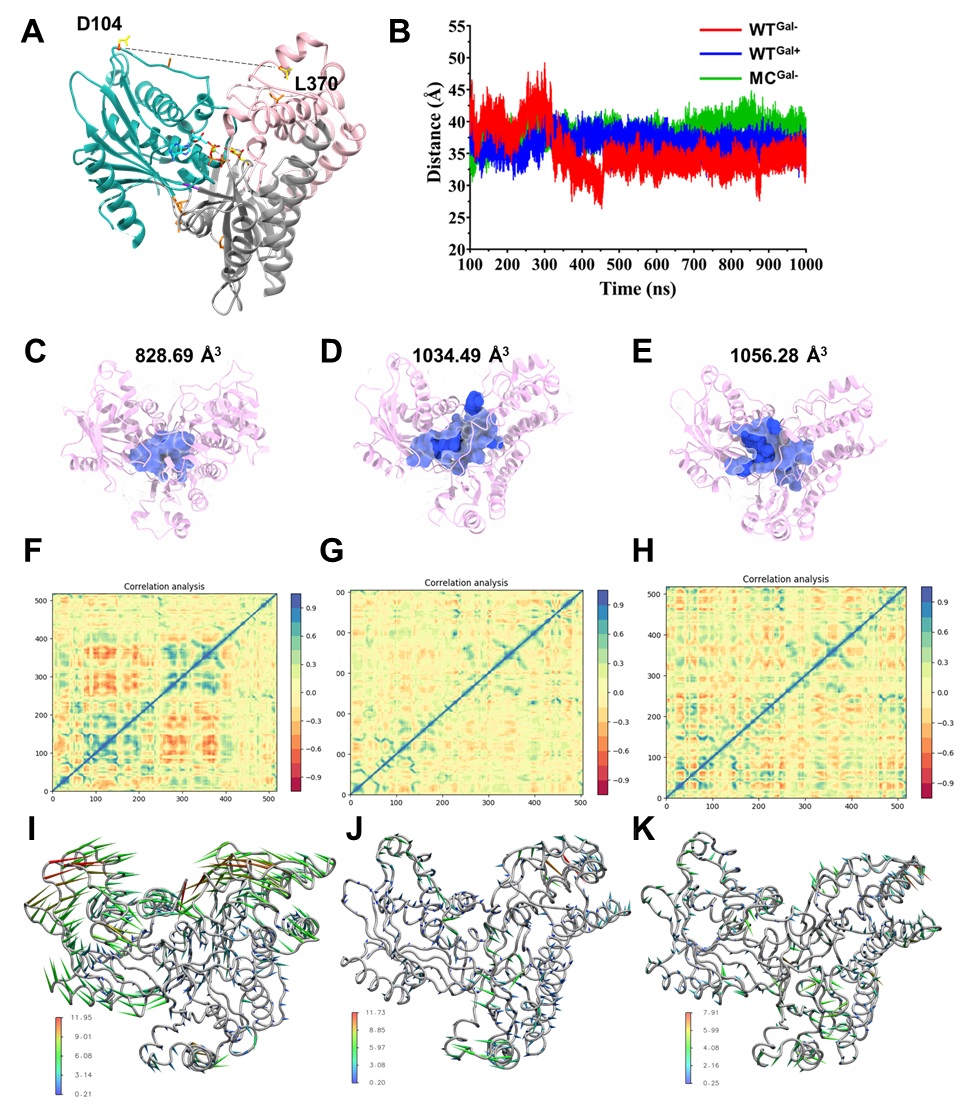

**Figure S5: Observations of 1000 ns scale molecular dynamics (MD) simulations.** **(A)** Structural representation of Gal3p in the closed conformation with dotted lines representing the distance between residues L370 and D104, found on the opposing ‘moving’ domains (cyan- and pink-colored regions), that is monitored during the simulation. **(B)** Time series of the distance measured between residues L370 and D104 records domain movement across 1000 ns of simulations. Volume of active site pocket (depicted as blue spheres) calculated for the last frame of the simulation for the **(D)** WT^Gal–^, **(E)** WT^Gal+^, and **(F)** MC^Gal–^ systems. Cross correlation studies were used to derive residue-residue correlation heatmaps for **(G)** WT^Gal–^, **(H)** WT^Gal+^, and **(I)** MC^Gal–^. Cross-correlation maps showed an intense residue-residue correlation between residues 90–250 and residues 250–350 in WT^Gal–^ that was not observed in the other two systems. Porcupine plots constructed from a principal component analysis (PCA) of the dynamics from the MD trajectories of **(D)** WT^Gal–^, **(E)** WT^Gal+^ and **(F)** MC^Gal–^ use spikes to show the intensity and direction of the principal component of motion for each residue. WT^Gal–^ showed approximately three-fold and four-fold greater motion relative to WT^Gal+^ and MC^Gal–^, respectively. Regions showing increased dynamics in the PCA of WT^Gal-^ correspond to the same regions showing defined residue-residue correlation from the cross-correlation studies.

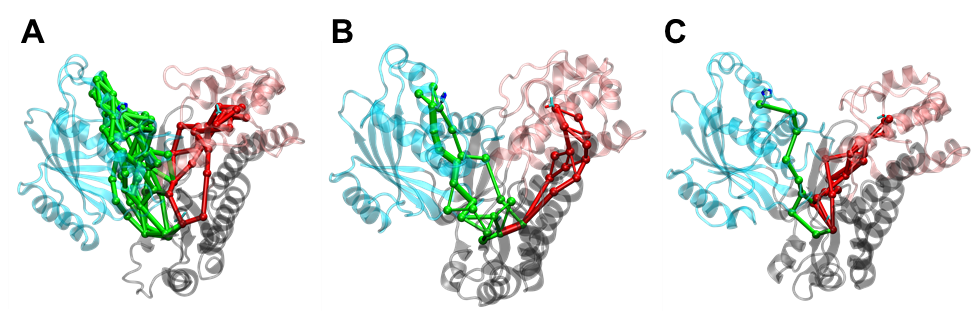

**Figure S6:** Residue-residue interaction networks were generated for the **(A)** WT^Gal–^, **(B)** WT^Gal+^, and **(C)** MC^Gal–^ systems using the NetworkView plugin in Visual Molecular Dynamics. Interaction networks between the static hinge (grey) region and the moving (cyan and pink) regions are visualized using green and red ball-and-stick models, respectively. Spheres represent nodes that are residues while edges represent the interaction between the residues with the thickness of the edge indicating the strength of interaction. The large number of interaction networks seen in WT^Gal–^ likely contributes to the formation the of compact, ‘collapsed’ structure observed during the molecular dynamics simulations. In contrast, the WT^Gal+^ and MC^Gal–^ systems exhibit much sparser interaction networks.

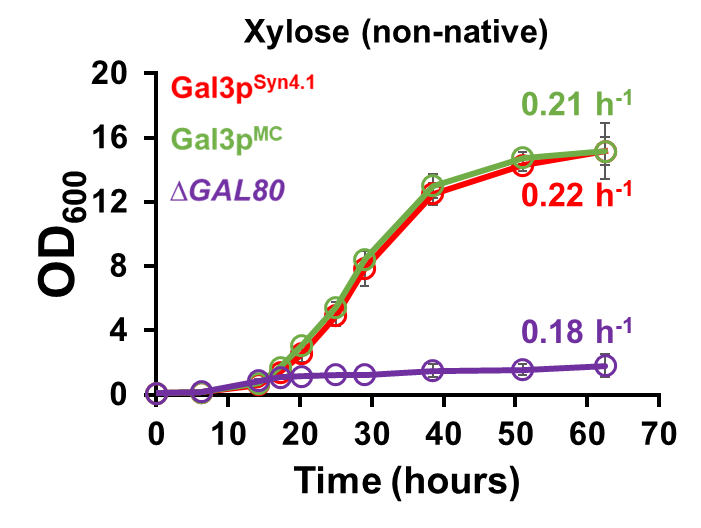

**Figure S7**: Comparing growth of the three GAL regulon activation systems on the non-native sugar xylose after being sub-cultured from glucose. Gal3p^Syn4.1^ requires induction by xylose, Gal3p^MC^ does not require an inducer but maintains the native regulatory architecture, while *∆GAL80* effects inducer-independent activation by removing native regulatory elements. All data points represent the average of three biological replicates ± sd.

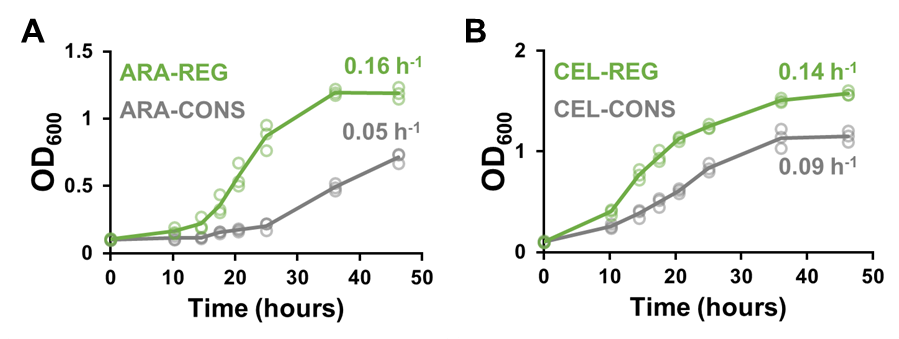

**Figure S8**: **Using Gal3p^MC^ to rapidly apply a regulon approach to growth on the non-native sugars, arabinose and cellobiose.** Growth of CONS and REG strains in defined media containing with in **(A)** 2 % arabinose or **(B)** 2 % cellobiose as the sole carbon source. All data points represent the average of three biological replicates ± sd.

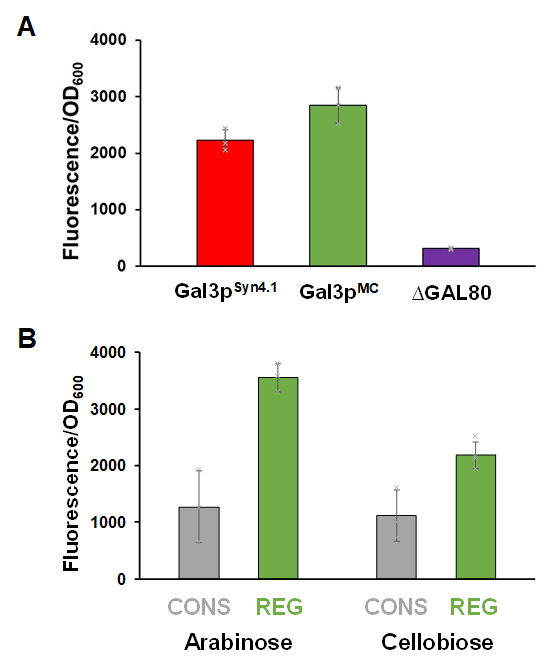

**Figure S9:** **Relative expression of catabolic genes in strains shown in Figure 5.** Cell-density normalized expression of a fluorescent protein (*EGFP*) from the representative promoters *GAL1p* and *TEF1p* used to control catabolic gene expression in REG and CONS strains, respectively, was quantified during exponential growth (t = 14 hours). These values serve as a proxy for the relative catabolic gene expression occurring in strains grown on xylose utilizing **(A)** three different approaches – substrate-specific (Gal3p^Syn4.1^), substrate-agnostic (Gal3p^MC^), and de-regulated (*∆GAL80*) GAL regulon activation. **(B)** CONS and REG approaches to growth on arabinose and cellobiose as sole carbon sources. Each data point represents the average of at least three biological replicates ± sd.

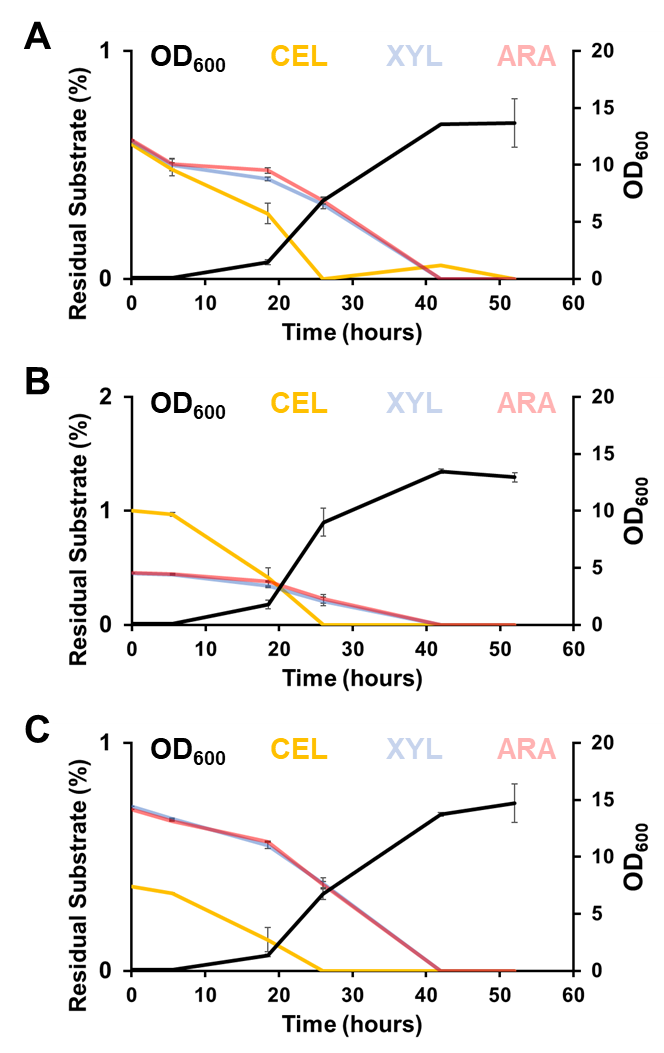

**Figure S10**: Cell density (black) and percent residual substrate of cellobiose (yellow), xylose (light blue), arabinose (pink), and both pentoses combined (purple) of Consolidated approach to co-utilization in complex media containing **(A)** 0.7 %-0.7 %-0.7 %, **(B)** 1 %-0.5 %-0.5 %, or **(C)** 0.4 %-0.8 %-0.8 % CEL-XYL-ARA (w/v). All data points represent the average of two biological replicates ± sd.

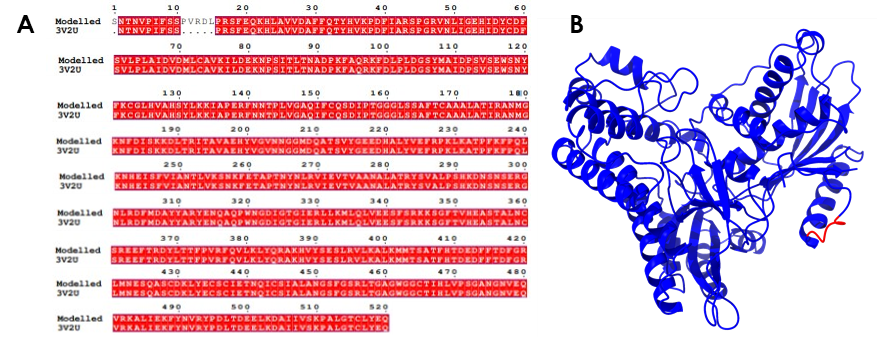

**Figure S11: (A)** Sequence alignment of the crystal structure (3V2U) of the closed state of Gal3p and the modelled structure of Gal3p protein. **(B)** 3D ribbon representation of the modelled structure of Gal3p protein, with the red loop indicating the modeled region spanning from residues 11 to 15 that was missing in the crystal structure.

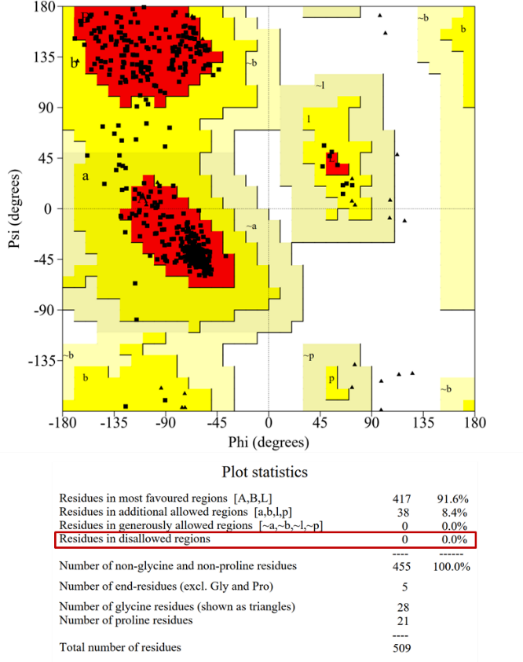

**Figure S12**: Ramachandran plot that was used to validate the model generated from SwissModel. This plot confirms that none of the residues in the model are in disallowed regions, indicating the high quality of our model.

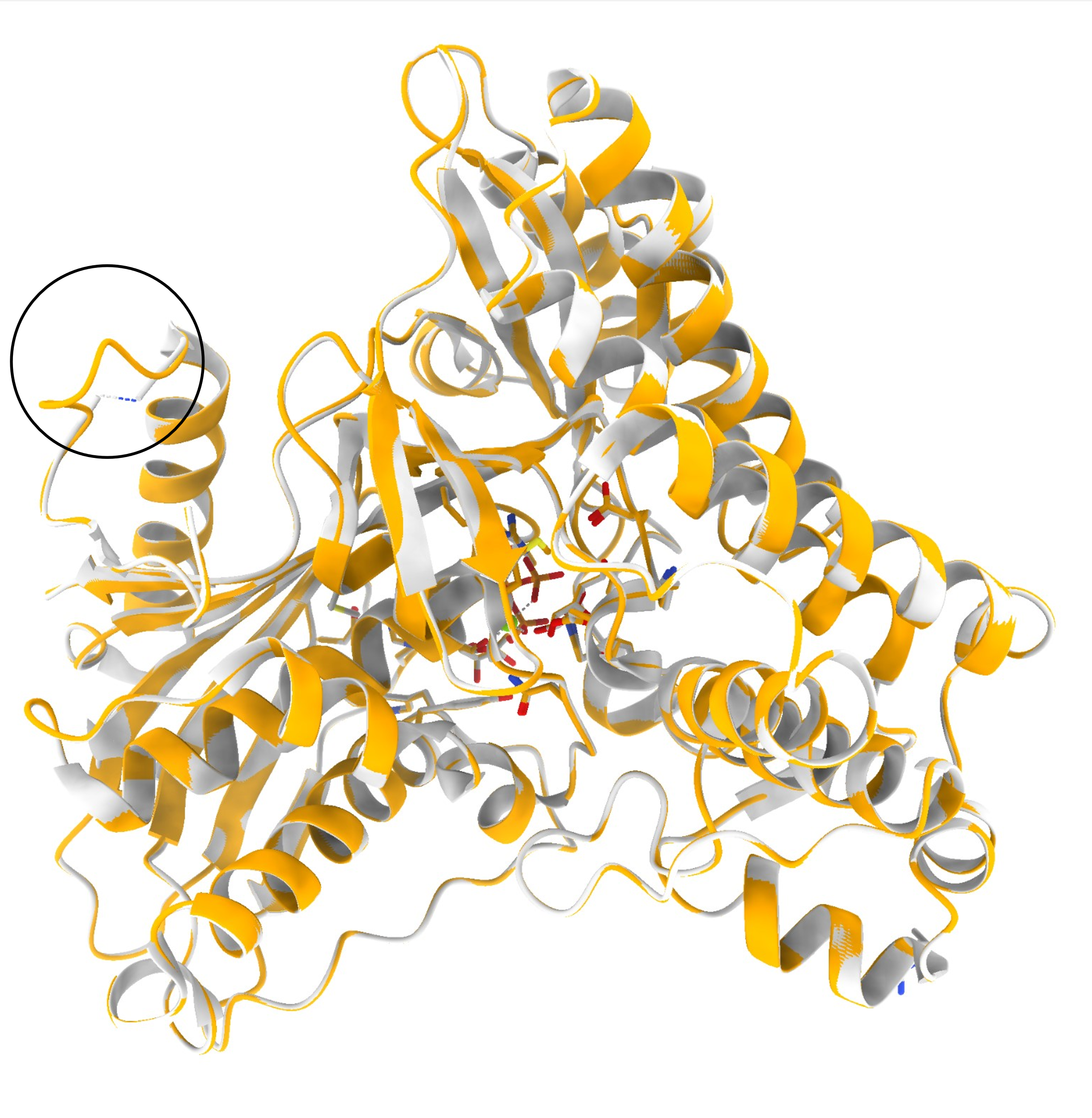

**Figure S13:** A comparative analysis was conducted to evaluate the accuracy of the models generated by Swiss-Model in relation to the closed-form crystal structure of Gal3p (PDB ID: 3V2U). The models were superimposed over the crystal structure in ribbon form, and the missing region in the model was highlighted. The Root Mean Square Deviation (RMSD) between the structures was calculated and found to be 0.09, indicating a close resemblance between the modelled and crystal structures.

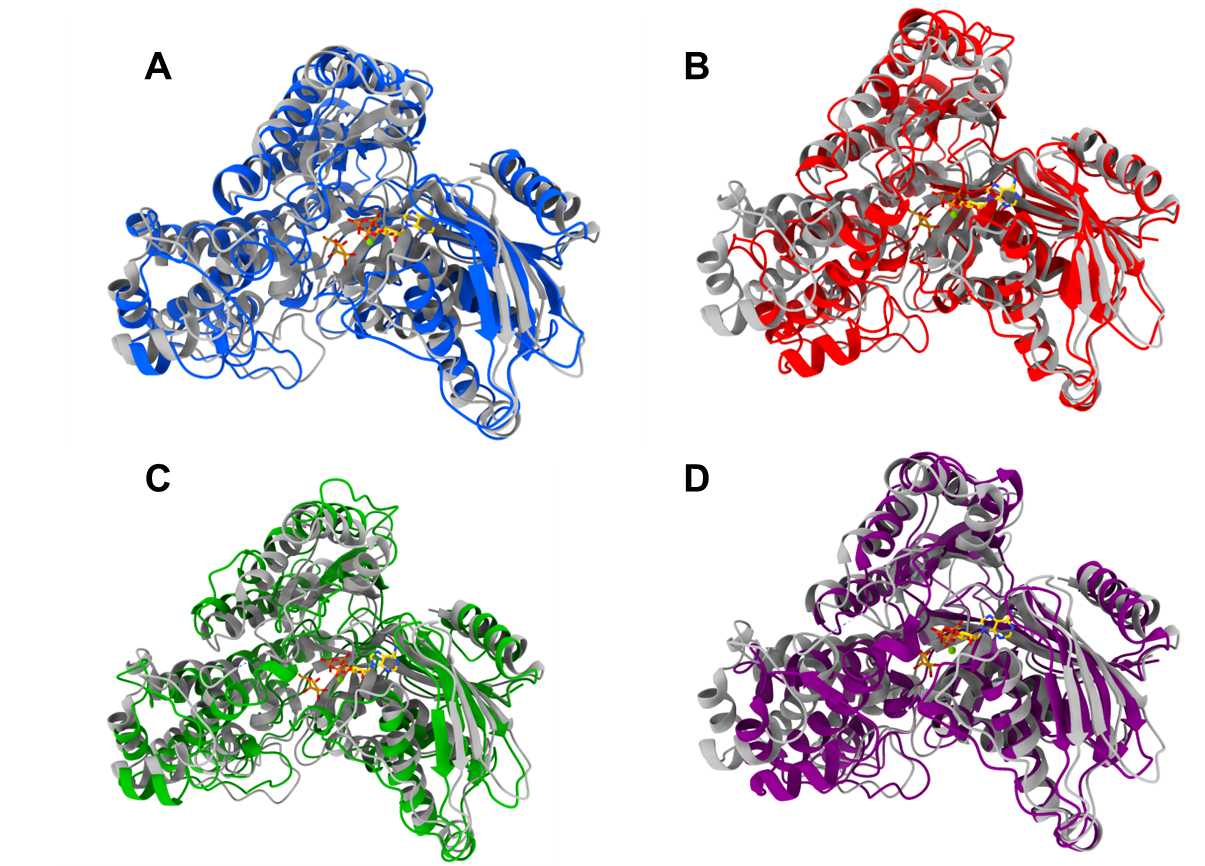

**Figure S14**: The superimposition analysis was conducted to compare the structures of Gal3p with the apo form crystal structure (grey ribbon, PDB ID 3V5R). The structures of WT^Gal+^ (blue), WT^Gal–^ (red), and MC^Gal–^ (green) were obtained from the minimum energy states of metadynamics simulations. The crystal structure of WT^Gal+^ in the closed state (violet, PDB ID 3V2U) was also included in the analysis. The RMSD values of the Cα backbone atoms were calculated to assess the structural differences. The RMSD values for WT^Gal+,^ WT^Gal–^, MC^Gal–^ with respect to the crystal structure in closed state were found to be 2.955, 4.422, and 3.220, respectively.
